# Tailored Design of Protein Nanoparticle Scaffolds for Multivalent Presentation of Viral Glycoprotein Antigens

**DOI:** 10.1101/2020.01.29.923862

**Authors:** George Ueda, Aleksandar Antanasijevic, Jorge A. Fallas, William Sheffler, Jeffrey Copps, Daniel Ellis, Geoffrey Hutchinson, Adam Moyer, Anila Yasmeen, Yaroslav Tsybovsky, Young-Jun Park, Matthew J. Bick, Banumathi Sankaran, Rebecca A. Gillespie, Philip J. M. Brouwer, Petrus H. Zwart, David Veesler, Masaru Kanekiyo, Barney S. Graham, Rogier Sanders, John P. Moore, Per Johan Klasse, Andrew B. Ward, Neil King, David Baker

**Author notes:** These authors contributed equally to this work (George Ueda, Aleksandar Antanasijevic). Correspondence to: Andrew Ward, Neil King, and David Baker.

## Abstract

The adaptive immune system is highly sensitive to arrayed antigens, and multivalent display of viral glycoproteins on symmetric scaffolds has been found to substantially increase the elicitation of antigen-specific antibodies. Motivated by the considerable promise of this strategy for next-generation anti-viral vaccines, we set out to design new self-assembling protein nanoparticles with geometries specifically tailored to scaffold ectodomains of different viral glycoproteins. We first designed and characterized homo-trimers from designed repeat proteins with N-terminal helices positioned to match the C termini of several viral glycoprotein trimers. Oligomers found to experimentally adopt the designed configuration were then used to generate nanoparticles with tetrahedral, octahedral, or icosahedral symmetry. Examples of all three target symmetries were experimentally validated by cryo-electron microscopy and several were assessed for their ability to display viral glycoproteins via genetic fusion. Electron microscopy and antibody binding experiments demonstrated that the designed nanoparticles display conformationally intact native-like HIV-1 Env, influenza hemagglutinin, and prefusion RSV F trimers in the predicted geometries. This work demonstrates that novel nanoparticle immunogens can be designed from the bottom up with atomic-level accuracy and provides a general strategy for precisely controlling epitope presentation and accessibility.

## Introduction

Multivalent antigen display, in which antigen is presented to the immune system in a repetitive array, has been demonstrated to increase the potency of humoral immune responses^1,2^. This has been attributed to increased cross-linking of antigen-specific B cell receptors at the cell surface and modulation of the immunogen trafficking to and within lymph nodes^3,4^. An ongoing challenge has been to develop multimerization platforms capable of presenting complex oligomeric or engineered antigens^5–7^, as these can be difficult to stably incorporate into non-protein-based nanomaterials (e.g. liposomes, polymers, transition metals and their oxides). Attributes such as epitope accessibility, proper folding of the antigen, and overall stability are additional considerations that must be taken into account in any design strategy for antigen presentation. Several reports have utilized non-viral, naturally occurring protein scaffolds, such as self-assembling ferritin^8–10^, lumazine synthase^5,11^, or encapsulin^12^ nanoparticles, to present a variety of complex oligomeric or engineered antigens. These studies collectively highlight several of the key advantages of using self-assembling proteins as scaffolds for multivalent antigen presentation^13,14^, including the formation of stable, highly monodisperse immunogens, the existence of scalable manufacturing methods, and seamless integration of antigen and scaffold via genetic fusion. More recently, computationally designed one-^15^ and two-^16,17^component protein nanoparticles have also been used to present complex oligomeric antigens^18,19^. The high stability of designed proteins^20,21^, versatility and ease of production and purification, increased potency observed upon immunization^18,19^, and the ability to predictively explore new structural space make these materials attractive as scaffolds for multivalent antigen presentation.

Despite this progress, control over the structure of nanoparticle immunogens has remained incomplete, limiting the potential of structure-based vaccine design. Even in the reported cases where computationally designed nanoparticles were used as scaffolds, the nanoparticle subunits were derived from naturally occurring oligomeric proteins. Nanoparticle scaffolds specifically designed from the bottom up for multivalent display of antigens of interest could be still more effective. For example, for scaffolding homo-oligomeric class I viral fusion proteins, a large group that includes many important vaccine antigens^22^, designed nanoparticles with a close geometric match between the C termini of the antigen and the N termini of the scaffold would provide considerable advantages. First, the antigen could be displayed without structural distortion near its base, potentially allowing for better retention of epitopes relevant to protection. Second, they could enable the multivalent display of antigens for which no compatible nanoparticle scaffolds are currently available. More broadly, by allowing atomically precise control over the geometry of antigen presentation, bottom up design of novel nanoparticle scaffolds would enable systematic investigation of the structural determinants of immunogenicity.

## Results

### Method overview

We sought to develop a general computational method for generating *de novo* protein nanoparticles with geometries tailored to display specific antigens of interest, focusing in particular on the challenge of displaying the pre-fusion conformations of the trimeric viral glycoproteins HIV-1 Env (BG505 SOSIP)^23,24^, influenza hemagglutinin (H1 HA)^25^, and respiratory syncytial virus (RSV) F (DS-Cav1)^7^. To make the tailored nanoparticle design problem computationally tractable, we used a hierarchical approach. First, monomeric protein building blocks were docked into trimeric homo-oligomers with cyclic symmetry (Figure 1a) and computationally screened for configurations featuring N termini with spacings similar to the C termini of the corresponding antigen (Figure 1b). Rosetta combinatorial sequence design was then used to generate energetically favorable protein-protein interfaces between the monomeric subunits. Upon experimental characterization, trimers found to self-assemble into the desired structure were then paired with second oligomeric components as building blocks for two-component nanoparticle docking in tetrahedral, octahedral, or icosahedral symmetry as previously described (Figure 1c)^16,17^. Rosetta sequence design was employed to optimize interactions between the oligomers, generating secondary designed interfaces to drive nanoparticle assembly. This step-wise computational design protocol yielded two-component nanoparticles tailored to present 4, 8, or 20 trimers of viral glycoprotein antigens (Figure 1d). These steps are described in detail in the following sections.

**Figure 1.**
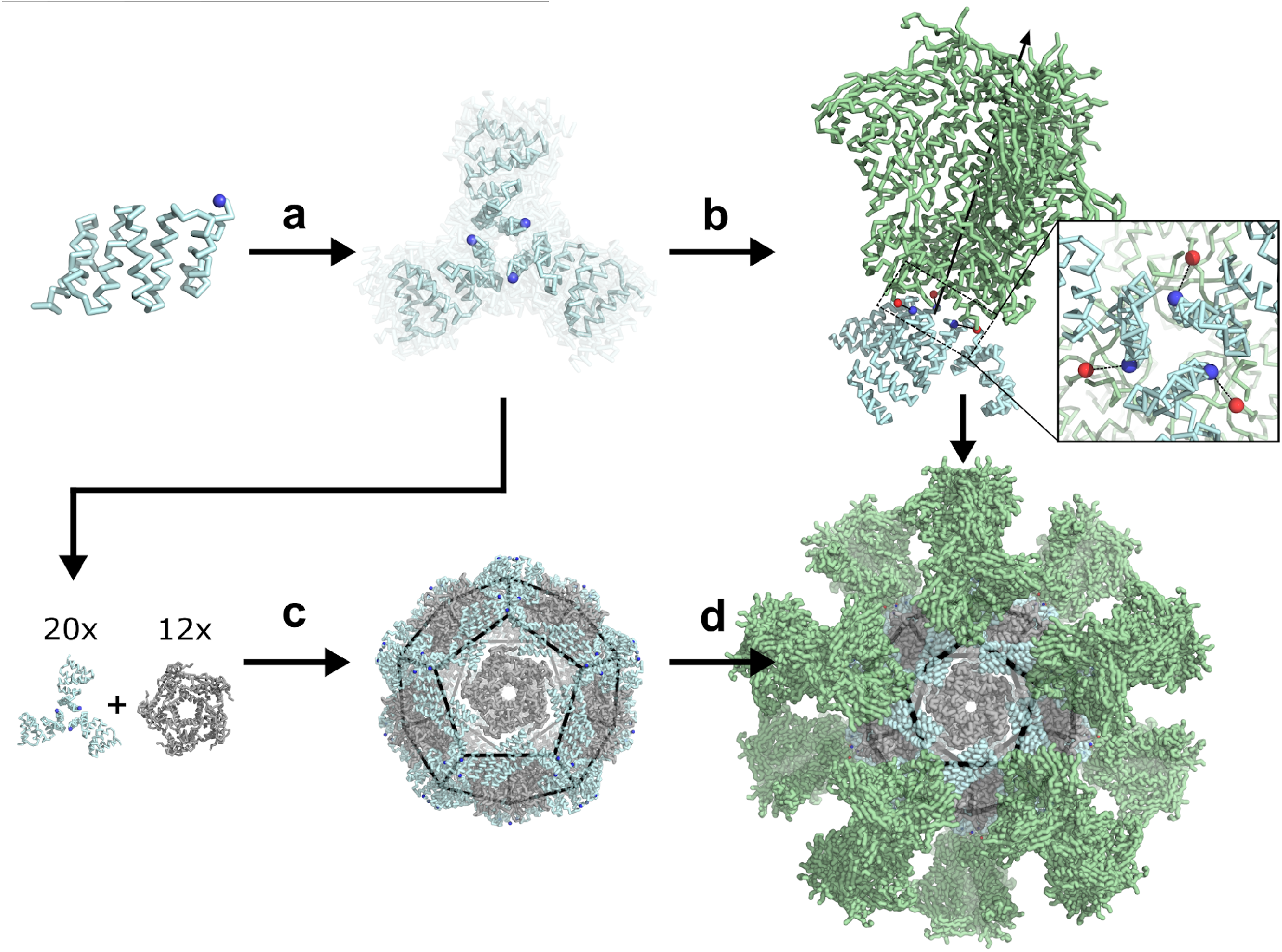
Hierarchical design of nanoparticles geometrically tailored to multivalently present specific oligomeric antigens. **a,** Computational docking and RPX scoring of C3-symmetric trimer docks from monomeric scaffold. **b,** Selection of trimer docks for Rosetta design and experimental screening based on termini alignment to antigenic target (green). **c,** Pairing of structurally-validated trimers with partner oligomers for two-component nanoparticle design. **d,** Replacement of designed trimeric component with antigen-fused counterpart yields multivalent nanoparticle.

### *In silico* symmetric docking and trimeric antigen targeting

A topologically diverse set of stable designed repeat proteins was used as scaffolds for symmetric trimer design, restricting trimeric configurations to those compatible with genetic fusion to at least one of the BG505 SOSIP, H1 HA, or DS-Cav1 trimers (see Methods). These viral fusion proteins were selected because of their importance as vaccine antigens, the availability of high-resolution structural data, and the variety they present in C-terminal geometry (31 nm, 15 nm, and 8 nm C-terminal separation distance for each glycoprotein, respectively). C3-symmetric docks of individual repeat proteins were rapidly assessed by the previously described RPX scoring method, which identifies arrangements likely to have good side chain packing^26^. Top-scoring docked configurations with an RPX score above 5.0 were computationally screened for termini geometries compatible with fusion to any of the three selected antigens using the previously described sic_axle protocol^18^. Geometrically compatible docks (those with non-clashing termini separation distances of 15 Å or less) were subjected to full Rosetta C3-symmetric interface design and filtering (see Methods)^26^, and 23 designs were selected for experimental characterization (design strategy presented in Supplementary Figure 1; sequences for all trimer designs are in Supplementary Table 7).

### Structural characterization of designed trimers

Synthetic genes encoding each of the designed trimers were expressed in *E. coli* and the resulting proteins purified from lysates by Ni^2+^ immobilized metal affinity chromatography (Ni^2+^ IMAC) followed by size-exclusion chromatography (SEC). 21 designs were found to express in the soluble fraction, and 11 formed the intended oligomeric state as assessed by SEC in tandem with multi-angle light scattering (SEC-MALS; examples in Figure 2 top panel, second row; SEC-MALS chromatograms for the remaining designs are in Supplementary Figure 2 and data in Supplementary Table 2; SEC chromatograms for remaining designs with off-target retention volumes are in Supplementary Figure 3). 4 of the designs that were trimeric and expressed well were selected for solution small angle X-ray scattering (SAXS) experiments and displayed scattering profiles very similar to those computed from the corresponding design model (Figure 2 top panel, third row; metrics in Supplementary Table 1 and 3), suggesting similar supramolecular arrangement, and these were carried forward into crystallization screens. Crystals were obtained for two designs, and structures of 1na0C3_2 and 3ltjC3_1v2 were determined at 2.6 and 2.3 Å resolution, revealing a heavy-atom R.M.S.D. between the design model and structure of each trimer at 1.4 and 0.8 Å, respectively. The structures confirmed in both cases that the designed proteins adopt the intended trimeric configurations, and that most of the atomic details at the trimeric interfaces are recapitulated as designed. Crystallization conditions, structure metrics, and structure-to-model comparisons are described in the Supplementary Methods, Supplementary Figure 4, and Supplementary Table 4.

**Figure 2.**
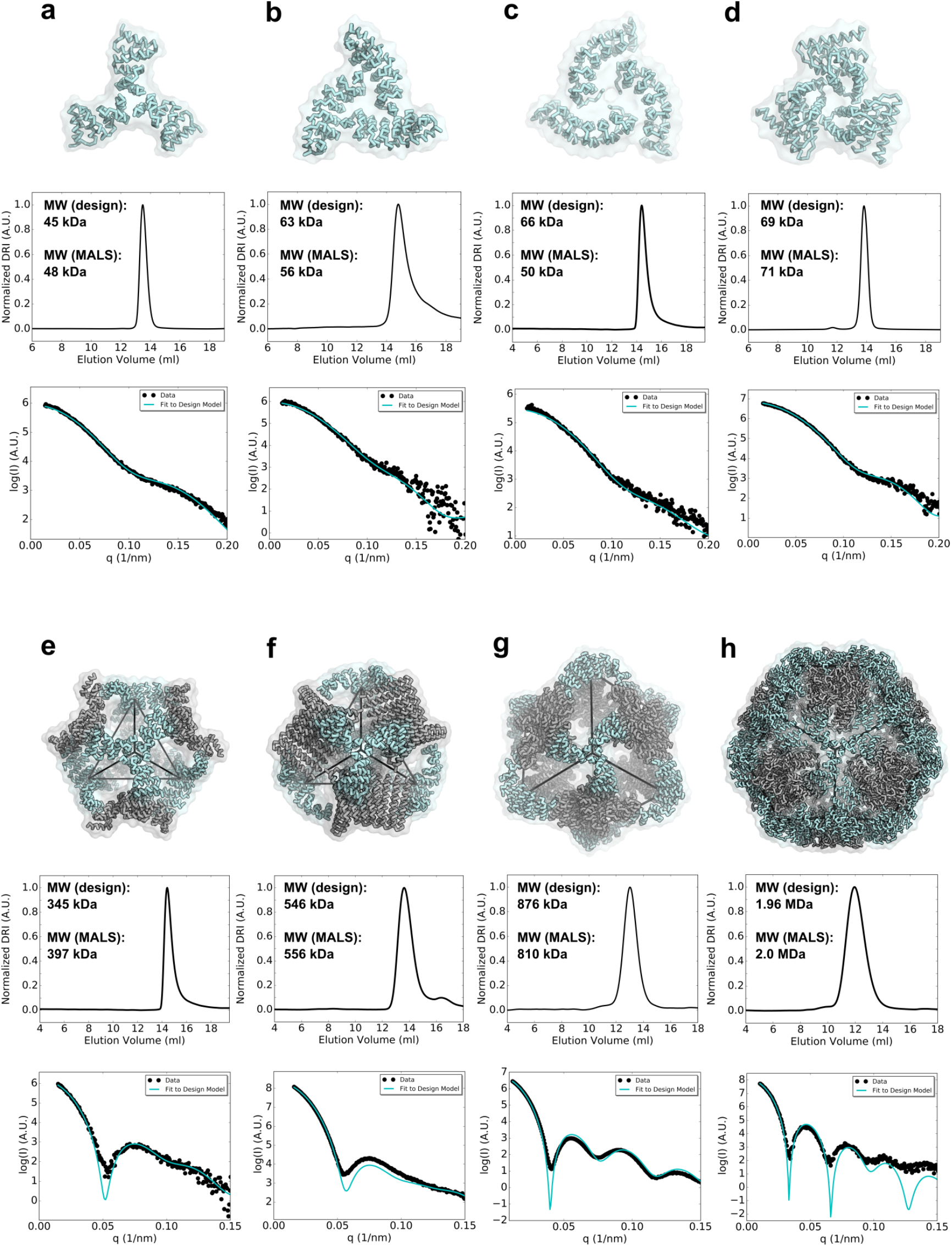
Biophysical characterization of antigen-tailored trimers and two-component nanoparticles. Top rows, design models. Middle rows, SEC chromatograms and calculated molecular weights from SEC-MALS. Bottom rows, SAXS comparisons between experimental data and curves calculated from design models. **a,**1na0C3_2. **b,**3ltjC3_1v2. **c,**3ltjC3_11. **d,** HR04C3_5v2. **e,** T33_dn2. **f,** T33_dn10. **g,** O43_dn18. **h,** I53_dn5.

### Generation of two-component nanoparticles from designed trimers

Each SAXS-validated trimer (Figure 2a-d) was docked pairwise with a set of previously designed symmetric homo-oligomers^26^ to generate tetrahedral, octahedral, and icosahedral arrangements using the TCdock program^16,17^. Icosahedral designs are particularly interesting as vaccine scaffolds as they have the highest valency among the cubic point group symmetries (20 trimers per particle compared to 8 for octahedra and 4 or 8 for tetrahedra), and higher valency has been shown to elicit higher antibody (Ab) titers in immunization studies^18,19,27^. To increase the probability of generating icosahedra, three naturally occurring homopentamers were also included in the docking calculations (PDB IDs 2JFB, 2OBX, and 2B98; sequences in Supplementary Table 7). To identify configurations likely to have a high number of energetically favorable side chain contacts following Rosetta interface design, nanoparticle docks were scored and ranked using the RPX method^26^. High-scoring and non-redundant nanoparticle configurations in which the N-terminal antigen fusion sites faced outward were selected for Rosetta interface design. The models with the best overall metrics for each unique docked configuration were then selected for experimental characterization (see Methods). The names of the 11 tetrahedra, 21 octahedra, and 21 icosahedra selected indicate, in order, the symmetry of the nanoparticle (T, O, or I), the oligomeric state of the first component (A), that of the second component (B), the letters “dn” reflecting the *de novo* origin of the input oligomers, and the rank by RPX score from the docking stage (e.g., “I53_dn5” indicates an icosahedral nanoparticle constructed from pentameric and trimeric components, ranked 5th in RPX-scoring). Sequences and input oligomer combinations for each designed nanoparticle are listed in Supplementary Table 8.

Synthetic genes encoding each of the 53 two-component nanoparticles were obtained and the designs were purified using Ni^2+^ IMAC (see Supplementary Methods). Pairs of proteins at the expected molecular weights were found to co-elute by SDS-PAGE for 24 of the designs, suggesting spontaneous association and pulldown of both components via only one His_6_-tagged component (exemplary co-eluting designs in Supplementary Figure 7). SEC chromatograms revealed that 19 designs did not form assemblies of the expected size or that the resulting assemblies were heterogeneous (Supplementary Figure 9). 5 designs, T33_dn2, T33_dn5, T33_dn10, O43_dn18, and I53_dn5, however, ran as monodisperse particles of the predicted molecular mass by SEC-MALS and were further investigated by SAXS. The experimentally determined solution scattering curves closely matched scattering curves computed from the design models^28^ for all 5 designs (Figure 2, bottom panel and Supplementary Figure 8; metrics in Supplementary Table 1 and 3, bottom 5 designs). Due to its high valency, the individual components of I53_dn5 were investigated for their capacity to be purified separately and assembled *in vitro*. Upon re-cloning, expression, and separate purification through a C-terminal His_6_-tag on each component, spontaneous nanoparticle assembly was achieved to near completion within minutes after equimolar mixing (Supplementary Figure 10).

### Structural Characterization of Designed Nanoparticles

The five SAXS-validated nanoparticles were structurally characterized using negative stain electron microscopy (NS-EM)^29,30^. 2,000–5,000 particles were manually picked from the electron micrographs acquired for each designed nanoparticle and classified in 2D using the Iterative MSA/MRA algorithm (see Supplementary Methods). 3D classification and refinement steps were performed in Relion/3.0^31^. Analysis of the NS-EM data confirmed high sample homogeneity for all 5 nanoparticle designs as evident from the micrographs and 2D class-averages (Figure 3). Free nanoparticle components were detected in the T33_dn5 sample, suggesting a certain propensity towards disassembly (Figure 3b). Analysis of the reconstructed 3D maps revealed that all 5 nanoparticles assemble as predicted by the design models, at least to the resolution limits of NS-EM.

**Figure 3.**
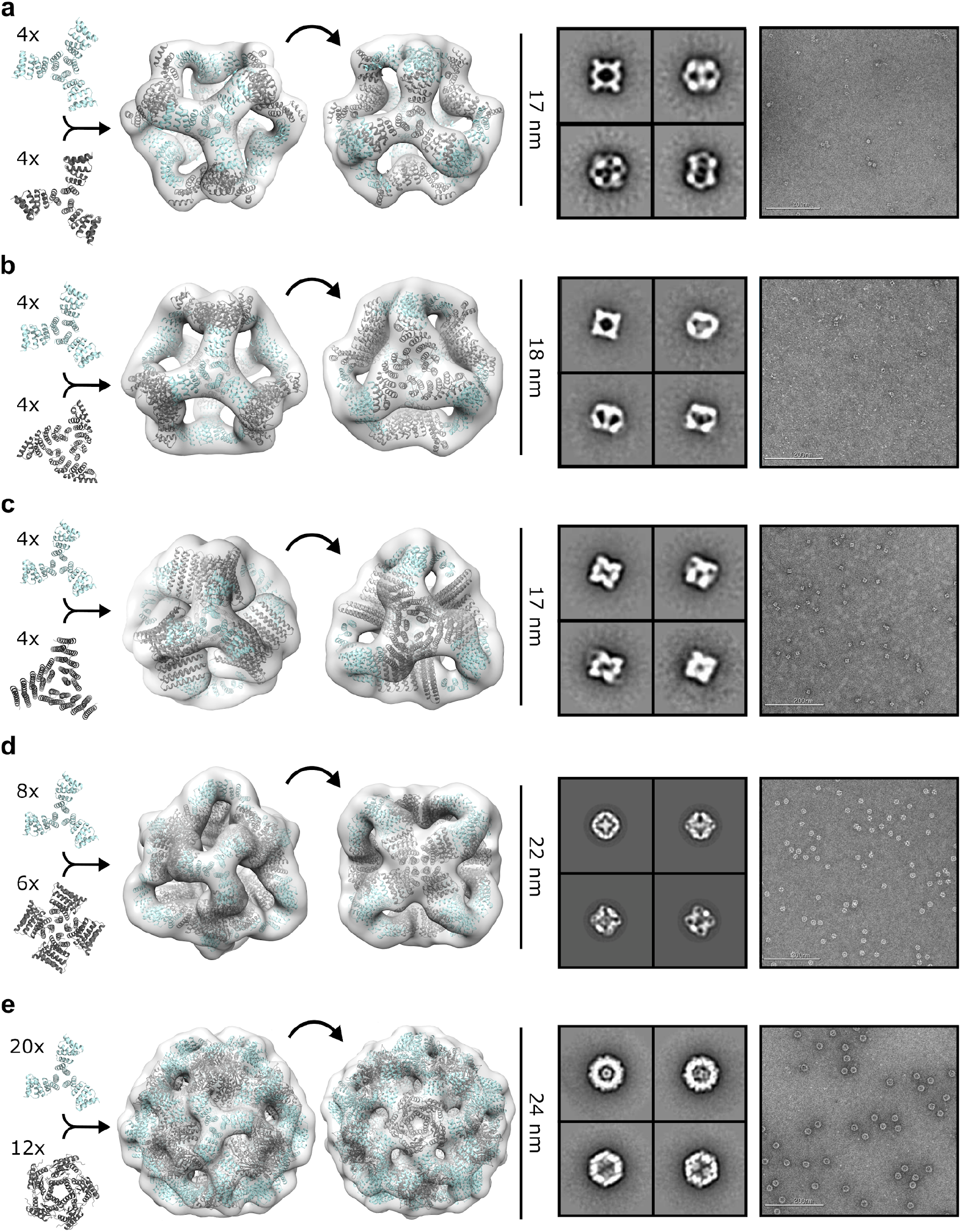
NS-EM analysis of antigen-tailored nanoparticles. From left to right: individual oligomeric components in nanoparticle assembly, design models fit into three-dimensional NS-EM reconstructions (views shown down each component’s axis of symmetry), 2D class-averages, raw electron micrographs of assembled particles. **a,** T33_dn2. **b,** T33_dn5. **c,** T33_dn10. **d,** O43_dn18. **e,** I53_dn5.

In order to obtain higher-resolution information, three nanoparticles, T33_dn10, O43_dn18, and I53_dn5, representing one of each targeted symmetry in the design stage (T, O, I), were subjected to cryo-electron microscopy (cryo-EM). Cryo-EM data acquisition was performed as described in the Methods section and data acquisition statistics are displayed in Supplementary Table 5. The data processing workflow is presented in Supplementary Figure 11. Appropriate symmetry (T, O, and I for T33_dn10, O43_dn18, and I53_dn5, respectively) was applied during 3D classification and refinement and maps were post-processed in Relion/3.0^31^. The final resolutions of the reconstructed maps for the T33_dn10, O43_dn18, and I53_dn5 nanoparticles were 3.9, 4.5, and 5.3 Å, respectively. Nanoparticle design models were relaxed into the corresponding EM maps by applying multiple rounds of Rosetta relaxed refinement^32^ and manual refinement in Coot^33^ to generate the final structures. Refined model statistics are shown in Supplementary Table 6. Reconstructed cryo-EM maps for T33_dn10, O43_dn18, and I53_dn5 and refined models are shown in Figure 4. Overall, the refined structures show excellent agreement with the corresponding Rosetta design models. Backbone R.M.S.D. values estimated for the asymmetric unit (consisting of a single subunit of component A and component B) were 0.65, 0.98, and 1.3 Å for T33_dn10, O43_dn18, and I53_dn5, respectively (Supplementary Table 1).

**Figure 4.**
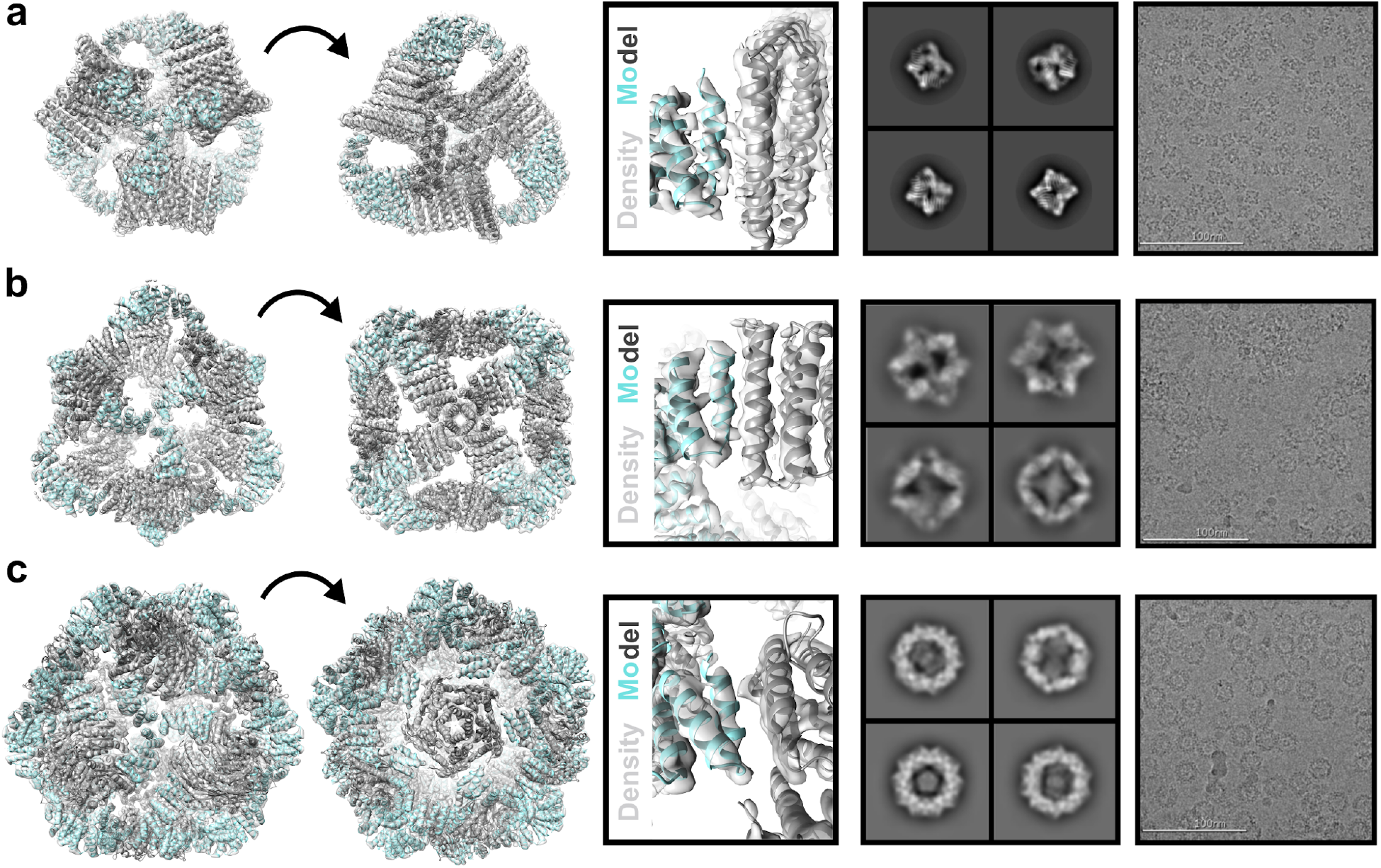
Cryo-EM analysis of antigen-tailored nanoparticles. From left to right: cryo-EM map with refined model fit into electron density, view of designed nanoparticle interface region fit into electron density, 2D class-averages, raw cryo-EM micrographs of assembled nanoparticles. **a,** T33_dn10. **b,** O43_dn18. **c,** I53_dn5.

### Characterization of Viral Glycoprotein-Nanoparticle Fusions

To explore the capability of the designed nanoparticles to display viral glycoproteins, we first generated fusions of a stabilized version of the BG505 SOSIP trimer to trimeric components of the nanoparticles. Synthetic genes for BG505 SOSIP fused to the N termini of T33_dn2A, T33_dn10A, and I53_dn5B (BG505 SOSIP– T33_dn2A, BG505 SOSIP–T33_dn10A, and BG505 SOSIP–I53_dn5), were transfected into HEK293F cells (see Supplementary Methods; sequences in Supplementary Table 9). The secreted fusion proteins were then purified using a combination of immuno-affinity chromatography and SEC. The second component for each nanoparticle was produced recombinantly in *E. coli* as described above for the designed trimers, and *in vitro* assembly reactions were prepared with equimolar mixtures of the two components and incubated overnight (see Supplementary Methods).

Assembled nanoparticles were purified by SEC and analyzed by NS-EM as described in the Methods section to assess particle assembly and homogeneity. ~1,000 particles were manually picked and used to perform 2D classification and 3D classification/refinement in Relion^31^. Reconstructed 3D maps with docked nanoparticle (cyan for fusion component and gray for assembly component) and BG505 SOSIP trimer (green) models are displayed in Figure 5 (left). BG505 SOSIP trimers are clearly discernible in 2D class-averages and reconstructed 3D maps. However, the trimers appear less well-resolved than the corresponding nanoparticle core in the three reconstructions, likely due to the short flexible linkers between the BG505 SOSIP trimer and the trimeric nanoparticle components (sequences in Supplementary Table 9). The self-assembling cores of the antigen-bearing T33_dn2, T33_dn10, and I53_dn5 nanoparticles were very similar to the NS-EM maps of the unmodified nanoparticles (at least to the resolution limits of NS-EM), demonstrating that fusion of the BG505 SOSIP trimer did not induce any major structural changes to the underlying nanoparticle scaffolds.

**Figure 5.**
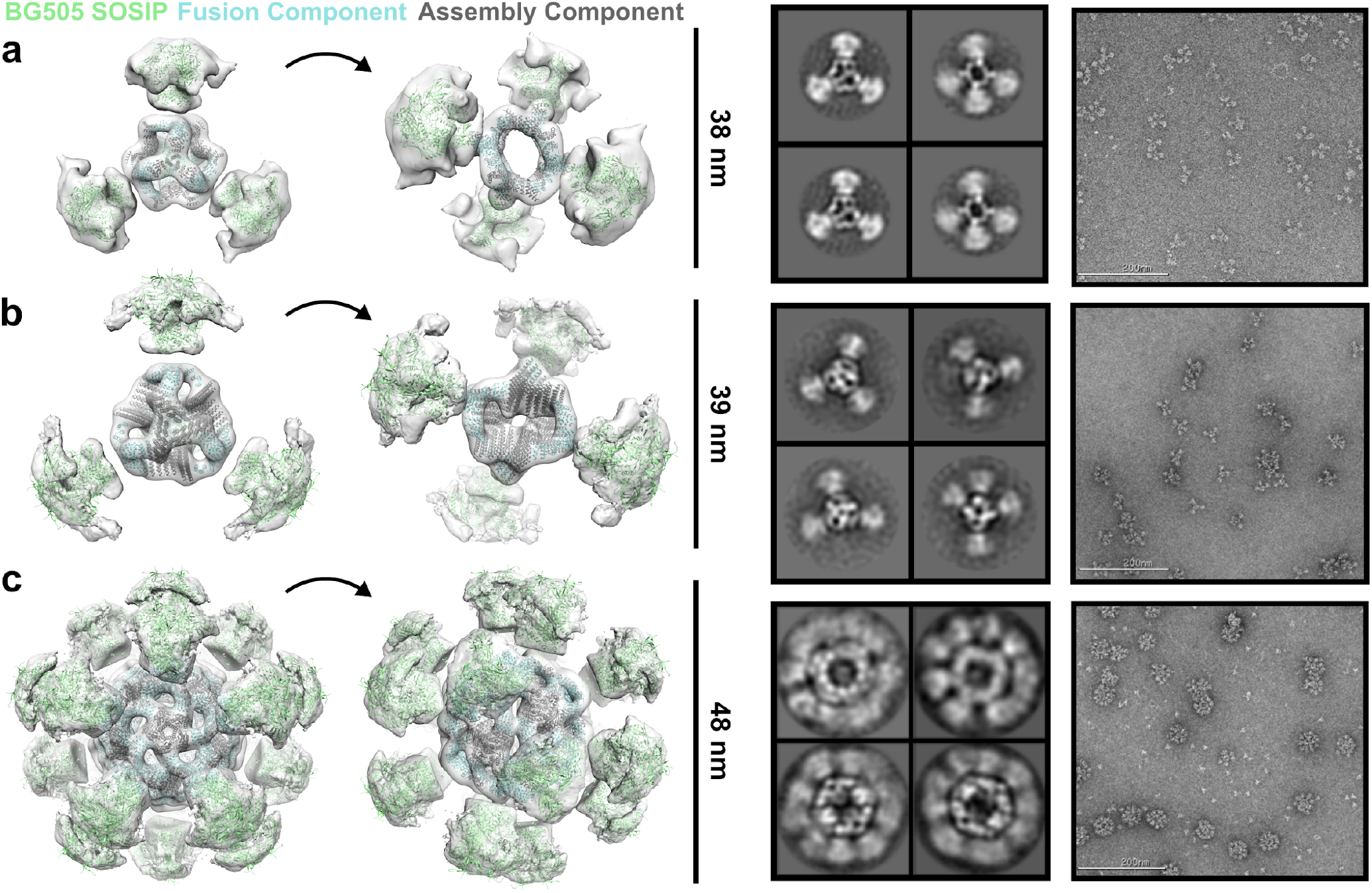
Structural validation of BG505 SOSIP antigen-fused particles. From left to right: nanoparticle design models and BG505 SOSIP trimer fit into 3D NS-EM reconstruction, 2D class-averages, raw NS-EM micrographs of assembled antigen-bearing nanoparticles. **a,** BG505 SOSIP–T33_dn2. **b,** BG505 SOSIP–T33_dn10. **c,** BG505 SOSIP–I53_dn5.

To further characterize the capability of the designed nanoparticles to display viral glycoproteins, we characterized the structures and antigenic profiles of I53_dn5 fused to the RSV F and influenza HA glycoproteins (DS-Cav1–I53_dn5 and HA–I53_dn5). Constructs were generated with the two viral glycoproteins fused to the N terminus of the I53_dn5B trimer (sequences in Supplementary Table 9), and the proteins were secreted from HEK293F cells and purified by Ni^2+^ IMAC. The fusion proteins were mixed with equimolar pentameric I53_dn5A, and the assembly reactions subjected to SEC. For both assemblies, the majority of the material migrated in the peak expected for assembled nanoparticles, and NS-EM analysis showed formation of I53_dn5-like nanoparticles with spikes emanating from the surface (Supplementary Figures 5 and 6). In both cases there was considerable variation in the spike geometry, again suggesting flexibility between the glycoproteins and the underlying particle. The GG linker connecting DS-Cav1 to I53_dn5 likely accounts for the observed flexibility in that case, which resulted in poor definition of the glycoprotein trimer in two-dimensional class averages (Supplementary Figure 5, bottom right). In HA–I53_dn5 there was no engineered linker between the glycoprotein and the nanoparticle trimer, and more clearly defined spike density was observed in the class averages (Supplementary Figure 6, bottom right).

**Figure 6.**
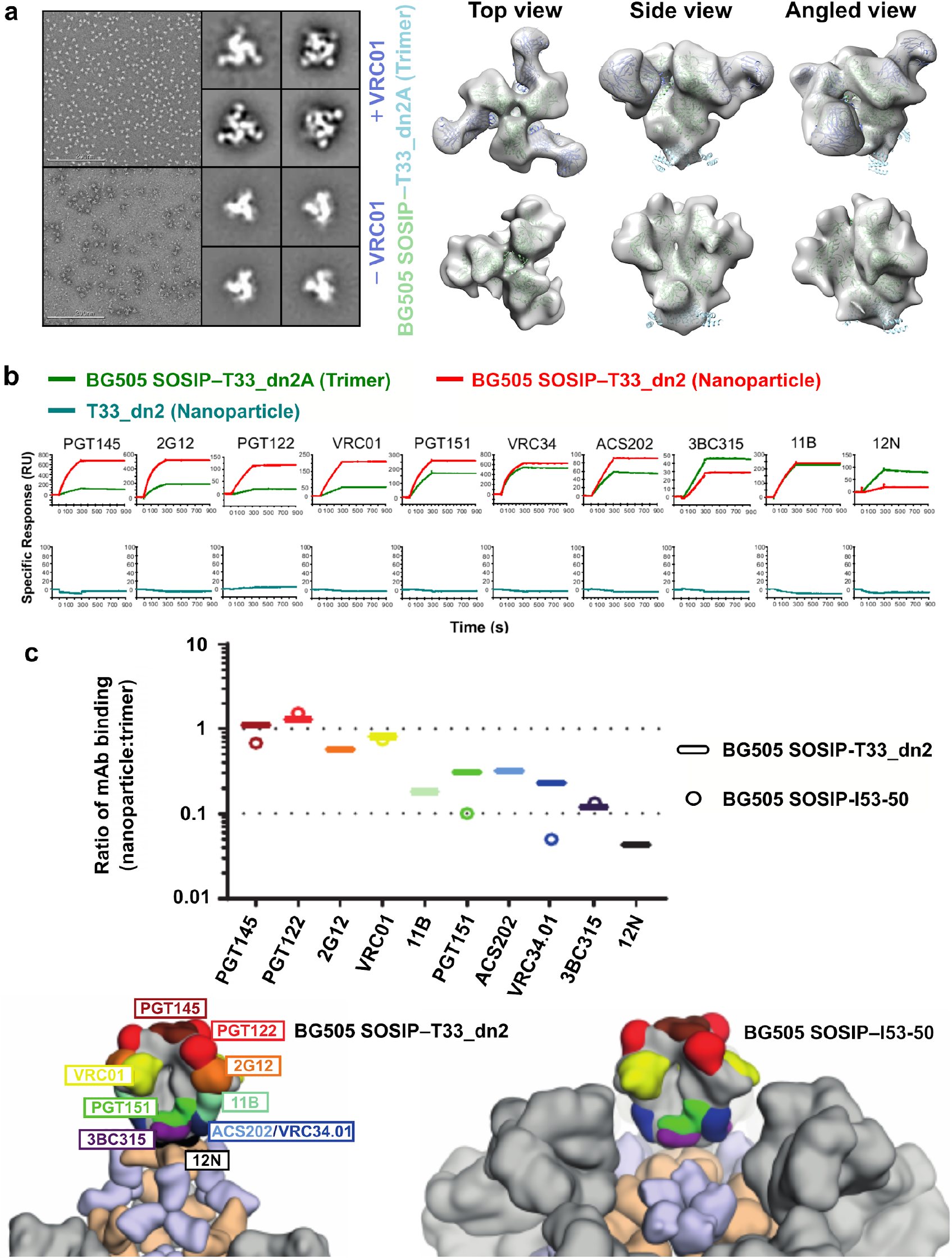
Tuning epitope accessibility through antigen display in monovalent, tetrahedral, and icosahedral formats. **a,** NS-EM micrographs of BG505 SOSIP–T33_dn2A with and without VRC01 Fab bound, 2D class averages, and models fit into 3D maps. **b,** Representative sensorgrams of indicated antigen binding to anti-Env mAbs. **c,** Relative accessibility of epitopes on BG505 SOSIP–T33_dn2 nanoparticles and BG505 SOSIP–I53-50 nanoparticles (reproduced from prior publication for comparison^19^) as determined by mAb binding (top, diagram). Ratio of moles of macromolecules are means of 2-4 experimental replicates. Epitopes mapped onto BG505 SOSIP are presented on either T33_dn2 or I53-50 (bottom, space-filling model). Wheat, antigen-bearing nanoparticle trimer component; purple, second nanoparticle component; gray, neighboring BG505 SOSIP trimers on the nanoparticle surface.

To determine if the displayed glycoproteins were properly folded, we examined their reactivity with conformationally specific monoclonal antibodies (mAbs). The DS-Cav1–I53_dn5 nanoparticle was found by ELISA to bind the RSV F-specific mAbs D25, Motavizumab, and AM14 similarly to soluble DS-Cav1 trimer with foldon^7^, indicating that the F protein is displayed in the desired prefusion conformation on the nanoparticle (Supplementary Figure 5, top). Biolayer interferometry binding experiments with anti-HA head^34^- and stem^35^-specific mAbs analogously showed that both the HA–I53_dn5 nanoparticle and the trimeric component HA–I53_dn5B presented the head and stem regions with wild-type–like antigenicity (Supplementary Figure 6, top).

### Control over BG505 SOSIP Epitope Accessibility using Different Display Geometries

Because previous work involving icosahedral nanoparticle scaffolds displaying HIV-1 Env trimers has shown that antigen crowding can modulate the accessibility of specific epitopes and thereby influence the humoral immune response^19,36^, we characterized the antigenicity of the BG505 SOSIP fusion nanoparticles in more detail. The designed nanoparticle scaffolds developed in this work exhibit varying geometries and valencies, providing a novel way to manipulate epitope accessibility. To determine the effect of particle symmetry/geometry on epitope accessibility, we selected the BG505 SOSIP–T33_dn2 nanoparticle as a candidate to compare against a previously published SOSIP-displaying icosahedral nanoparticle (BG505 SOSIP–I53-50)^19^ through surface plasmon resonance (SPR) experiments. To first validate mAb binding to the BG505 SOSIP trimer on a structural level when the latter is fused to the trimeric component T33_dn2A, NS-EM class averages and a 3D reconstruction (see Methods) were carried out with the VRC01 Fab, and the previously observed binding mode was confirmed (Figure 6a). Next, a panel of anti-Env mAbs targeting epitopes ranging from the apex to the base of the BG505 SOSIP trimer were immobilized on SPR sensor chips (Figure 6b). BG505 SOSIP–T33_dn2A trimer or BG505 SOSIP–T33-dn2 nanoparticle was then flowed over the mAbs and the ratio of macromolecule bound was calculated from the binding signal as previously described^19^. For mAbs that target apical, V3-base, and CD4-binding site epitopes (PGT145, PGT122, 2G12, and VRC01), the number of molecules of trimer or nanoparticle bound was relatively similar (ratio ~1). However, for mAbs that target more base-proximate epitopes in the gp120-gp41 interface (ACS202, VRC34, and PGT151), an inter-protomeric gp41 epitope (3BC315), and the main autologous neutralizing Ab epitope in the glycan hole centered on residues 241 and 289 (11B), the accessibility was reduced in the nanoparticle format. Binding to a mAb directed to the trimer base (12N) was obliterated on nanoparticle BG505 SOSIP–T33_dn2 (Figure 6b). We also compared this binding to historic data for BG505 SOSIP–I53-50 using six different representative mAbs^19^. As for BG505 SOSIP–T33_dn2, mAbs to the apex, V3-base, and CD4-binding site (PGT145, PGT122, and VRC01) gave molar ratios ~1 for BG505 SOSIP–I53-50. However, for mAbs that target the more base-proximate epitopes in the gp120-gp41 interface (VRC34 and PGT151), there was a nearly 3-fold higher epitope accessibility on T33_dn2 than I53-50 nanoparticles (Figure 6c). Further down the trimer, no accessibility difference was again observed for a mAb that targets the gp41 inter-protomeric epitope (3BC315), which was relatively inaccessible on both nanoparticles, likely due to steric hindrance by neighboring trimers. These findings demonstrate the ability to modulate epitope accessibility by varying the designed scaffold geometry; this capability conferred by the panel of designed nanoparticles could be extremely useful in structure-based approaches to eliciting epitope-targeted humoral immune responses.

## Discussion

Strong BCR signaling is required for eliciting successful humoral immune responses, but the molecular mechanisms by which this can be accomplished are not fully understood. Historically, live-attenuated or inactivated viruses and engineered virus-like particles (VLPs) have been able to confer protective immunity without pathogenicity, but the empirical discovery and compositional complexity of such vaccines has hampered understanding of possible mechanisms for eliciting protective levels of antibodies. Antigen display on multivalent carriers has become a general strategy for enhancing antigen-specific Ab titers by orders of magnitude^1,2^. The new generation of antigen-tailored protein nanoparticles introduced in this study provides a systematic way to optimize immune responses by varying the geometry of antigen presentation.

Success in the *de novo* design of homo-trimers and subsequent two-component nanoparticles showcases the capabilities of computational protein design to generate structures on the nanometer scale from scratch. The repeat proteins used here as building blocks—many of which were themselves designed *de novo*—are exceptionally stable, conferring stability to the nanoparticles which they form. The design of the trimeric units to be structurally compatible with viral glycoprotein antigens in a targeted manner is a new accomplishment. These constructs likely stabilize their fused antigens as their termini were purposefully tethered to match the conformations found in previously published crystal structures^23,37,38^. This study demonstrates, both at the oligomer and nanoparticle design stages, the capability of generating protein-based materials across a range of controllable sizes and topologies for targeted biological applications. These proteins represent a panel of molecules well-suited to probe not only the adaptive immune response, but also other biological systems across various size scales.

The ability to fully customize the structures of nanoparticle scaffolds could be particularly useful for HIV-1 Env-based immunogens. While previous studies of nanoparticle-displayed HIV-1 Env trimers have demonstrated enhanced immunogenicity^39^, the effects are often modest compared to those observed for other antigens^1,2,18,36^. While there may exist intrinsic peculiarities to HIV-1 Env that limit increases in Ab responses upon multivalent display^40,41^, limitations associated with epitope inaccessibility caused by crowding of the trimers on nanoparticle surfaces have also been identified^19^. This observation strongly motivates the exploration of antigen presentation geometry through the custom design of nanoparticle scaffolds to most effectively elicit the desired immune response. Indeed, the SPR experiments presented here demonstrate that epitopes proximate to the BG505 SOSIP base were markedly more accessible to immobilized mAbs on BG505 SOSIP–T33_dn2 than BG505 SOSIP–I53-50. Furthermore, the availability of multiple antigen-displaying nanoparticles makes possible the usage of different nanoparticle scaffolds during prime and boost immunizations, which could limit immune responses directed toward the nanoparticle scaffolds while boosting antigen-specific Ab responses. Finally, the flexibility in the valency and geometry of antigen presentation enabled by bottom up nanoparticle design provides a route to investigate the effects of these parameters on B cell activation and the potency and breadth of the ensuing humoral response. This capability could help overcome the intrinsically low affinity of germline BCRs for fusion glycoproteins, thus enabling bNAb development that would otherwise not occur.

## Methods

### Targeted Docking and Trimer Design Against Selected Antigens

The set of scaffolds were used for symmetric trimer design targeted against three viral fusion protein topologies: HIV-1 BG505 SOSIP, influenza H1 HA, and RSV F protein (PDB IDs 5VJ6 res. 518-664, 5KAQ res. 11-501, 5TPN res. 27-509)^23,37,38^. A C3-symmetric specified docking search was first performed, and output was assessed by the previously described RPX scoring method for symmetric docking^26^. Up to the top-scoring 100 docks for each scaffold were aligned against each of the three immunogens along the shared axis of symmetry and sampled translationally along and rotationally about the axis in 1 Å and 1° increments, respectively. For each sample, the distance between the C terminal residue of the target immunogen and N terminal residue of the docked trimer was measured until a minimum, non-clashing distance was determined (Supplemental Figure 1). Solutions for docks that were less than or equal to 15 Å for one or more of the three immunogens were selected for full Rosetta symmetric interface design as described in previously published methods^26^. Individual design trajectories were filtered by the following criteria: difference between Rosetta energy of bound (oligomeric) and unbound (monomeric) states less than −30.0 Rosetta energy units, interface surface area greater than 700 Å^2^, Rosetta shape complementarity (sc) greater than 0.65, and less than 50 mutations made from the respective native scaffold. Designs that passed these criteria were manually inspected and refined by single point reversions, and one design per unique docked configuration was added to the set of trimers selected for experimental validation.

### Computational Design of Nanoparticles Validated Trimers

Two-component nanoparticle docks were scored and ranked using the RPX method^26^ as opposed to prior methods involving only interface residue contact count^16^. High-scoring and non-redundant nanoparticle configurations were selected for Rosetta interface design with an added caveat that they included trimers with outward-facing N termini for antigen display. The design protocol took a single-chain input .pdb of each component, and a symmetry definition file^42^ containing information for a specified cubic point group symmetry. The oligomers were then aligned to the corresponding symmetry axes of the architecture using the Rosetta SymDofMover, taking into account the rigid body translations and rotations retrieved from the .dok file output from TCdock^16,17^. Each configuration was then subjected to a symmetric interface design protocol similar to that previously described^26^. Individual design trajectories were filtered by the following criteria: difference between Rosetta energy of bound and unbound states less than −30.0 Rosetta energy units, interface surface area greater than 700 Å^2^, sc greater than 0.6, and less than 50 mutations made from each native scaffold. Designs that passed these criteria were manually inspected and refined by single point reversions for mutations that did not appear to contribute to stabilizing the bound state of the interface. The sequence with the best overall metrics for each unique docked configuration was then selected for experimental characterization.

### NS-EM of T33_dn2, T33_dn5, T33_dn10, O43_dn18, I53_dn5, BG505 SOSIP–T33_dn2, BG505 SOSIP–T33_dn10, and BG505 SOSIP–I53_dn5

NS-EM experiments were performed as described previously^29,30^. Fusion components and assembled nanoparticle samples (with and without antigen) were diluted to 20-50 μg/ml and loaded onto the carbon-coated 400-mesh Cu grid that had previously been glow-discharged at 15 mA for 25 s. VRC01 complexes with BG505 SOSIP–T33_dn2A were formed by combining the trimer with a 6-fold molar excess of the VRC01 Fab and subsequent incubation for 1 hour at room temperature. Complex sample was diluted to 20 μg/ml and loaded onto the glow discharged Cu grids. Grids were negatively stained with 2% (w/v) uranyl-formate for 60 s. Data collection was performed on a Tecnai Spirit electron microscope operating at 120 keV. The magnification was 52,000× with a pixel size of 2.05 Å at the specimen plane. The electron dose was set to 25 e^−^/Å^2^. All imaging was performed with a defocus value of −1.50 μm. The micrographs were recorded on a Tietz 4k×4k TemCam-F416 CMOS camera using Leginon automated imaging interface. Data processing was performed in Appion data processing suite. For BG505 SOSIP-fused nanoparticle samples (termed v5.2(7S), see Supplementary methods), approximately 500–1000 particles were manually picked from the micrographs and 2D-classified using the Iterative MSA/MRA algorithm. For non-fused nanoparticle samples, 2,000–5,000 particles were manually picked and processed. For BG505 SOSIP-fused trimer samples and Fab complexes, 10,000–40,000 particles were auto-picked and 2D-classified using the Iterative MSA/MRA algorithm. 3D classification and refinement steps were done in Relion/2.1^43^. The resulting EM maps have been deposited to EMDB with IDs: 21162 (T33_dn2), 21163 (T33_dn5), 21164 (T33_dn10), 21165 (O43_dn18), 21166 (I53_dn5), 21167 (BG505 SOSIP–T33_dn2A), 21168 (BG505 SOSIP–T33_dn2A + VRC01 Fab), 21169 (BG505 SOSIP–T33_dn2 nanoparticle), 21170 (BG505 SOSIP–T33_dn10 nanoparticle), 21171 (BG505 SOSIP–I53_dn5 nanoparticle).

### NS-EM of DS-Cav1–I53_dn5

The sample was diluted with a buffer containing 10 mM HEPES pH 7.0 and 150 mM NaCl to a concentration of 0.025 mg/ml and adsorbed for 15 s to a glow-discharged carbon-coated copper grid. The grid was washed with the same buffer and stained with 0.7% uranyl formate. Images were collected at a nominal magnification of 57,000 using EPU software on a ThermoFisher Talos F200C electron microscope equipped with a 4k×4k Ceta camera and operated at 200 kV. The pixel size was 0.253 nm. Particles were picked automatically using in-house written software (unpublished) and extracted into 200×200-pixel boxes. Reference-free 2D classifications were performed using Relion 1.4^31^ and SPIDER^44^.

### NS-EM of HA–I53_dn5

The HA–I53_dn5 complex was adsorbed onto glow-discharged carbon-coated copper mesh grids for 60 s, stained with 2% uranyl formate for 30 s, and allowed to air dry. Grids were imaged using the FEI Tecnai Spirit 120 kV electron microscope equipped with a Gatan Ultrascan 4000 CCD Camera. The pixel size at the specimen level was 1.60 Å. Data collection was performed using Leginon^45^ with the majority of the data processing carried out in Appion^46^. The parameters of the contrast transfer function (CTF) were estimated using CTFFIND4^47^. All particles were picked in a reference-free manner using DoG Picker^48^. The HA–I53_dn5 particle stack from the micrographs collected was pre-processed in Relion. Reference-free two-dimensional (2D) classification with cryoSPARC was used to select a subset of particles, which were used to generate an initial model using the Ab-Initio reconstruction function in CryoSPARC. The particles from the best class were used for non-uniform refinement in CryoSPARC to obtain the final 3D reconstruction.

### Cryo-EM of Designed Nanoparticles

Grids were prepared on Vitrobot mark IV (Thermo Fisher Scientific). Temperature was set to 10 °C, humidity at 100%, wait time at 10 s, while the blotting time was varied in the 4-7 s range with the blotting force at 0. The concentrations of T33_dn10, O43_dn18, and I53_dn5 nanoparticle samples were 4.2, 3.0, and 1.9 mg/ml, respectively. n-Dodecyl β-D-maltoside (DDM) at a final concentration of 0.06 mM was used for sample freezing. Quantifoil R 2/1 holey carbon copper grids (Cu 400 mesh) were pre-treated with Ar/O_2_ plasma (Solarus plasma cleaner, Gatan) for 10 s prior to sample application. Concentrated nanoparticle samples were mixed with appropriate volumes of stock DDM solution and 3 μl applied onto the grid. Excess sample and buffer was blotted off and the grids were plunge-frozen into nitrogen-cooled liquid ethane. Cryo-grids were loaded into a Titan Krios (FEI) operating at 300 kV, equipped with the K2 direct electron detector camera (Gatan). Exposure magnification of 29,000 was set with the resulting pixel size of 1.03 Å at the specimen plane. Total dose was set to ~50 e^−^/Å^2^ with 250 ms frames. Nominal defocus range was set to −0.6 to −1.6 μm for all 3 nanoparticle samples. Automated data collection was performed using Leginon software^45^. Data collection information for acquired datasets is shown in Supplementary Table 5.

### Cryo-EM Data Processing

MotionCor2^49^ was run to align and dose-weight the movie micrographs. GCTF v1.06 was applied to estimate the CTF parameters. Initial processing was performed in cryoSPARC 2.9.0. Template-picked particles were extracted and subjected to 2D classification. Multiple rounds of heterogeneous refinement were performed to further clean-up particle stacks in 3 acquired datasets. Selected subsets of particles were then transferred to Relion 3.0^31^ for further processing. Reference models were generated using Ab-Initio reconstruction in cryoSPARC v2.9.0^50^ with the application of appropriate symmetry (tetrahedral, octahedral, and icosahedral for T33_dn10, O43_dn18, and I53_dn5, respectively). Several rounds of 3D classification and refinement were used to sort out a subpopulation of particles that went into the final 3D reconstructions. Tetrahedral, octahedral, and icosahedral symmetry restraints were applied for all 3D refinement and classification steps during the processing of T33_dn10, O43_dn18, and I53_dn5 datasets, respectively. Soft solvent mask around the nanoparticle core was introduced during the final 3D classification, refinement, and post-processing. The resolutions of the final reconstructed maps were 3.86 Å for T33_dn10, 4.54 Å for O43_dn18, and 5.35 Å for I53_dn5. The resulting EM maps have been deposited to EMDB with IDs: 21172 (T33_dn10), 21173 (O43_dn18) and 21174 (I53_dn5). A graphical summary of the data processing approach and relevant information for each dataset are displayed in Supplementary Figure 11.

### Surface Plasmon Resonance (SPR) Analysis of BG505 SOSIP Trimer and Nanoparticle Binding to Immobilized Broadly Neutralizing Abs (bNAbs)

The antigenicity of BG505 SOSIP–T33_dn2A trimer and BG505 SOSIP–T33_dn2 nanoparticle was analyzed on a BIAcore 3000 instrument at 25 °C and with HBS-EP (GE healthcare Life sciences) as running buffer, as described^19^. Briefly, affinity-purified goat anti-human IgG Fc (Bethyl Laboratories, Inc.) and goat anti-rabbit IgG Fc (Abcam) were amine-coupled to CM3 chips and human and rabbit mAbs were captured to an average density of 320 ± 1.5 RU (s.e.m). BG505 SOSIP–T33_dn2A or BG505 SOSIP–T33_dn2 (both v5.2(7S) without N241/N289, see Supplementary Methods and Supplementary Table 9) was allowed to associate for 300 s and then dissociate for 600 s at a concentration of 10 nM assembled macromolecule (i.e., trimer or nanoparticle). The low background binding in parallel flow cells with only anti-Fc was subtracted. The lack of binding of nanoparticles lacking Env was ascertained for each Ab. To illustrate how epitope accessibility affects the relative binding of the trimers and nanoparticles, we converted the signals, which are proportional to mass bound, to moles bound and calculated the ratio for nanoparticles/trimers. For this comparison historic data on icosahedral nanoparticles were included^19^. The number of moles binding to the immobilized IgG at the end of the association phase was calculated: 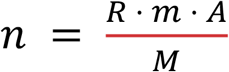where *n* is the number of moles of macromolecules, *R* the response at 300 s (RU), *m* the mass bound per area and RU (g/(mm^2^ RU)), *A* the interactive area of the chip (mm^2^), and *M* the molar mass of the macromolecule (g/mol). This analysis corrects for the greater mass (and thereby greater signal) for each bound nanoparticle such that the number of binding events by differing macromolecules can be directly compared.

## Acknowledgements

This work was supported by the Howard Hughes Medical Institute, the Bill and Melinda Gates Foundation (OPP1120319, OPP1111923, OPP1156262, OPP1115782 and OPP1084519), the National Science Foundation (NSF CHE 1629214), and the Intramural Research Program of the Vaccine Research Center, NIAID, NIH. We thank Lauren Carter at the Institute for Protein Design for assistance with SEC-MALS. We thank M. Capel, K. Rajashankar, N. Sukumar, J. Schuermann, I. Kourinov and F. Murphy at NECAT supported by grants from the National Center for Research Resources (5P41RR015301-10) and the National Institute of General Medical Sciences (P41 GM103403-10) from the National Institutes of Health. We thank Kathryn Burnett and Greg Hura for SAXS data collection through the SIBYLS mail-in SAXS program. This work was conducted at the Advanced Light Source (ALS), a national user facility operated by Lawrence Berkeley National Laboratory on behalf of the Department of Energy, Office of Basic Energy Sciences, through the Integrated Diffraction Analysis Technologies (IDAT) program, supported by DOE Office of Biological and Environmental Research. Additional support comes from the National Institute of Health project ALS-ENABLE (P30 GM124169) and a High-End Instrumentation Grant S10OD018483. X-ray crystallography data were collected at the ALS, and the Berkeley Center for Structural Biology is supported in part by the National Institutes of Health, National Institute of General Medical Sciences, and the Howard Hughes Medical Institute. The ALS is supported by the Director, Office of Science, Office of Basic Energy Sciences, of the U.S. Department of Energy under Contract No. DE-AC02-05CH11231.

## Structure Deposition Information

Crystal structures have been deposited in the RCSB protein data bank with the PDB IDs 6V8E (1na0C3_2) and 6VEH (3ltjC3_1v2). Cryo-EM structures have been deposited to the RCSB database with PDB IDs: 6VFH (T33_dn10), 6FVI (O43_dn18), and 6VFJ (I53_dn5). Electron density maps have been deposited in the EMDB with 21162 (T33_dn2), 21163 (T33_dn5), 21164 (T33_dn10), 231165 (O43_dn18), 21166 (I53_dn5), 21167 (BG505 SOSIP–T33_dn2A), 21168 (BG505 SOSIP–T33_dn2A + VRC01 Fab), 21169 (BG505 SOSIP–T33_dn2 nanoparticle), 21170 (BG505 SOSIP–T33_dn10 nanoparticle), 21171 (BG505 SOSIP–I53_dn5 nanoparticle), 21172 (T33_dn10, cryo-EM map), 21173 (O43_dn18, cryo-EM map), and 21174 (I53_dn5, cryo-EM map).

## Author contributions

G.U., A.A., J.A.F., A.W., N.K., and D.B. designed the research. G.U. carried out computational docking, antigen alignments, design calculations, and biophysically characterized designed trimers and nanoparticles. A.A. performed cryo-EM analysis on designed nanoparticles, optimized production of all BG505 SOSIP fusions, and performed NS-EM analysis of BG505 SOSIP fusion particles. J.A.F. and W.S. implemented RPX code into the Rosetta software suite used for trimer and nanoparticle design. J.C. performed additional NS-EM on designed nanoparticles. G.U. crystallized the designed trimers, M.J.B., B.S., and P.Z. processed the diffraction data and solved the crystal structures. D.E., Y.J.P., and D.V. produced and characterized HA–I53_dn5. G.H., R.A.G., Y.T., M.K., and B.S.G. produced and characterized DS-Cav1–I53_dn5. A.M. assisted in nanoparticle design. P.B., A.Y., J.M., and P.J.K. performed SPR experiments and quantitative comparison between BG505 SOSIP nanoparticles. All authors discussed results and commented on the manuscript.

