## Supplementary Information for "Tailored Design of Protein Nanoparticle Scaffolds for Multivalent Presentation of Viral Glycoprotein Antigens"

### Supplementary Methods

**Repeat Protein Scaffolds**. Repeat proteins, comprised of recurring 20-50 residue stretches, are ideal for use in protein-based material design due to their high stability and capability to have altered lengths and curvatures by varying the number of repeating modules. Listed below are the RCSB Protein Data Bank entries for scaffolds used in this study. An additional set of scaffolds is provided in which experimental small-angle X-ray scattering data agreed with the computational model[1].

| X-ray Structures | (PDB ID) | SAXS Validated Models |
|---|---|---|
| 1na0 | (1NA0) | tpr1 |
| 3ltj | (3LTJ) | HR00 |
| 2fo7 | (2FO7) | |
| HR04 | (5CWB) | |
| HR07 | (5CWD) | |
| HR10 | (5CWG) | |

**Protein Expression and Purification from E. coli.** Synthetic genes for designed proteins were optimized for *E. coli* expression and assembled from purchased genes (Genscript or Gen9) ligated into the pET21-NESG (designed trimers) or pET28b-NESG (designed nanoparticles) vector at restriction sites NdeI and XhoI or NcoI and XhoI respectively. A second ribosome-binding site was inserted between the open-reading frames of individual components of nanoparticle designs (“AGAAGGAGATATCAT”), such that the two proteins would be co-expressed and screened for co-elution by SDS-PAGE. Plasmids were cloned into BL21 or Lemo21 (DE3) *E. coli* competent cells. Transformants were inoculated and grown in 5 mL of either LB or TB medium with either 100 mg/L carbenicillin or 100 µg/L kanamycin at 37 °C overnight. Subsequently, liquid cultures were inoculated 1:100 (v:v) and grown at 37 °C until an $OD_{600}$ of 0.5-0.8. Isopropyl-thio-β-Dgalactopyranoside (IPTG) was then added at a concentration of 0.5-1 mM and growth temperature was reduced to 18 °C to induce protein expression, or cultures were left at 37 °C and auto-induced by virtue of the media-included galactose according to the Studier protocols[2]. Expression proceeded for 20 hours until the cell cultures were harvested by centrifugation. Cell pellets were resuspended in 25 mM Tris, 150 mM NaCl, 5mM imidazole, DNase, EDTA-free protease inhibitors (Pierce), pH 8.0. and lysed by sonication or microfluidization. Each protein was then purified from lysate by $Ni^{2+}$ immobilized metal affinity chromatography (IMAC) with Ni-NTA Superflow resin (Qiagen or GE). Resin with bound cell lysate was washed with 15 column volumes of 25 mM Tris, 150 mM NaCl, 40 mM imidazole, pH 8.0. Proteins were eluted with 5 column volumes of 25 mM Tris, 150 mM NaCl, 400 mM imidazole, pH 8.0 for further purification by size exclusion chromatography.

**Size-Exclusion Chromatography.** Elution samples for each designed protein were concentrated down using a 10,000 MWCO protein concentrator (Novagen) and fractionated by size on an AKTA pure chromatography system using a Superdex 200 or Superose 6 10/300 GL gel filtration column (GE Life Sciences) in 25 mM Tris, 150 mM NaCl, pH 8.0 (TBS). Sizing profiles were noted based on absorption at 220 nm and 280 nm wavelength light for each fraction. Molecular weights for predominant species in each protein trace were estimated by comparison to the corresponding monomeric profile.

**Analytical Size-Exclusion Chromatography on Sephacryl S-500.** Purified HA-bearing nanoparticles or trimers were applied to a Sephacryl S-500 column pre-equilibrated with

25 mM Tris, 2 M NaCl, 5% glycerol, pH 8.0 . Sizing profiles were recorded based on absorption at 280 nm wavelength light.

**Size-Exclusion Chromatography with Multi-Angle Light Scattering.** Fractions containing single predominant species from the initial round of size exclusion chromatography were concentrated down with 10,000 MWCO protein concentrators (Novagen) to a concentration of 1.0-2.0 mg/mL and run through a high-performance liquid chromatography system (Agilent) using a Superdex 200 or Superose 6 10/300 GL gel filtration column (GE Life Sciences) in TBS buffer. These fractionation runs were coupled to a multi-angle light scattering detector (Wyatt) in order to determine the absolute molecular weights for each designed protein complex.

**Small-Angle X-ray Scattering.** Designed proteins that predominantly formed the target oligomeric species were re-expressed and purified for low-resolution solution structure determination by small-angle X-ray scattering (SAXS) at the SIBYLS High Throughput SAXS Advanced Light Source in Berkeley, California[3]. A beam exposure time of between 0.3-10 seconds was used to obtain averaged diffraction data (SAXS FrameSlice Application), which are represented in plots of log intensity (I) vs. q. A 11kEV/1.125A X-ray beam was used with a 2 m beamstop.

**Crystallization Conditions.** Design 1na0C3_2 was found to crystallize in 1 M LiCl, 100 mM citrate, 20% w/v PEG 6000, pH 4, and was frozen using 25% glycerol as cryoprotectant. Design 3ltjC3_1 crystallized in 1 mM DL-glutamic acid monohydrate, 100 mM DL-alanine, 100 mM glycine, 100 mM DL-lysine monohydrochloride, 100 mM DL-serine; 100 mM Tris, 100 mM BICINE, 20% v/v ethylene glycol, 10% w/v PEG 8000, pH 8.5. Diffraction data for each of these designs were collected at the Advanced Light Source (Beamline 8.2.1) at Lawrence Berkeley National Laboratory in Berkeley, California. Both trimer designs contained uncleaved C terminal $His_6$-tags in crystallized conditions.

**Data collection, structure determination and refinement**. Diffraction data for 3ltjC3_1 was collected on beamline 5.0.1 at the Advanced Light Source (Berkeley, CA) and 1na0C3_2 on beamline 8.2.1, both using an ADSC Q315R CCD area detector. Both datasets were scaled and merged in HKL2000[4]. The structures were phased by molecular replacement, with the computational design as the search model, using the program PHASER[5] in the PHENIX software suite[6]. Iterative rounds of manual model building and refinement were conducted in Coot[7] and Phenix.refine[8], respectively for both structures. Hydrogens were added for all refinement runs. The geometric quality of the final model was assessed using the Molprobity server[9]. Resolution cutoff was determined by monitoring the refinement statistics in the context of the reflection data completeness and the $CC^{1/2}$ and $I/\sigma I$ values[10].

**I53_dn5 *in vitro* Assembly.** The ability of the two nanoparticle components of I53_dn5 to be separately purified and then mixed to achieve nanoparticle assembly was assessed. Genes for the individual oligomeric components were cloned into expression vectors and expressed independently in *E. coli*. The $His_6$-tagged proteins were purified following the purification protocol described above for the designed trimers. Initial size-exclusion chromatograms for the components were obtained on a Superdex 200 10/300 GL column, and predominant peak species were stoichiometrically mixed in TBS buffer for 20 minutes at 25 °C. A secondary size-exclusion step was performed on a Superose 6 10/300GL column to assess assembly of the intended particle based on expected retention volume.

**Nanoparticle Structural Model Building and Refinement from CryoEM data.** Post-processed maps from Relion were used to relax and refine nanoparticle models. Rosetta relaxed refinement[11] was performed for all datasets using the Rosetta design models of T33_dn10, O43_dn18, and I53_dn5 nanoparticles as inputs. Appropriate symmetry was enforced during model refinement. EMRinger[12] and Molprobity[12,13] scores were used to evaluate the output structures. The best models for each nanoparticle were then manually inspected and edited in Coot v0.9-pre[7]. For T33_dn10 and O43_dn18 datasets this procedure was repeated 2 more times to generate the final structures. The resulting models have been deposited to the PDB database with the IDs: 6VFH (T33_dn10), 6VFI (O43_dn18), and 6VFJ (I53_dn5). Model refinement statistics is shown in Supplementary Table 5.

**Production and Purification of BG505 SOSIP–T33_dn2A, BG505 SOSIP–T33_dn10A, and BG505 SOSIP–I53_dn5B.** Synthetic genes were optimized for mammalian expression and subcloned into pPPI4 vector. BamHI and NheI restriction sites were used for insertion of different nanoparticle assembly components to the C-terminus of BG505 SOSIP. Quick Ligation kit, BamHI-HF, and NheI-HF restriction enzymes were purchased from New England Biolabs (NEB). BG505 SOSIP variant used for all early optimizations steps was engineered with a combination of v5.2[14] (mutations: E64K, A73C, A316W, A561C) and MD3D[14,15] (mutations: M271I, A318Y, R585H, L568D, V570H, R304V, F519S) stabilizing mutations, and had glycosylation sites introduced at positions 241 and 289 (mutations: P240T, S241N, F288L, T290E, P291S). This construct was termed BG505 SOSIP.v5.2(7S). For epitope-accessibility experiments (by surface plasmon resonance), a version of this construct was designed without the 241 and 289 glycans. Protein sequences of different constructs used here are shown in Supplementary Table 9. HEK 293F cells were grown in suspension using FreeStyle 293 Expression Medium (Thermo Fisher Scientific) at 135 RPM, 8% $CO_2$, 80% humidity, 37 °C. At confluency of ~$1\times10^{-6}$ cells/ml, the cells were co-transfected with pPPI4 DNA vectors encoding the appropriate fusion component (250 µg per 1 L of cells) and furin protease (80 µg per 1 L of cells). Polyethyleneimine (Polysciences Inc) used as transfection reagent (1 mg per 1 L of cells). Cells were incubated for 6 days, after which they were spun down by centrifugation (7,000 RPM, 1 hour, 4 C) and the protein-containing supernatant was further clarified by vacuum-filtration (0.45 µm, Millipore Sigma). For immuno-affinity chromatography steps, Sepharose 4B columns with immobilized PGT145 IgG were used. Fusion components were eluted with 3 M magnesium chloride, 250 mM L-Arginine buffer, pH 7.2 into an equal volume of gel filtration buffer (25 mM Tris, 250 mM L-Arginine, 500 mM NaCl, 5 % glycerol, pH 7.4). Eluates were concentrated and buffer exchanged into gel filtration buffer. A Sephacryl S200 16/600 column was used for subsequent gel filtration.

**Production and Purification of HA–I53_dn5B.** Synthetic genes were optimized for mammalian expression and subcloned into the CMV/R vector (VRC 8400)[16]. XbaI and AvrII restriction sites were used for insertion of I53_dn5B component to the C terminus of the H1 HA ectodomain (residues 1-676 from A/Michigan/45/2015) which also contained a Y98F mutation to prevent sialic-acid binding and self-aggregation during expression[17]. Gene synthesis and cloning was performed by Genscript. The protein sequence of HA–I53_dn5B is shown in Supplementary Table 8. HEK 293F cells were grown in suspension using Expi293 Expression Medium (Thermo Fisher Scientific) at 150 RPM, 8% $CO_2$, 70% humidity, 37 °C. At confluency of ~$2.5\times10^{-6}$ cells/mL, the cells were co-transfected with the vector encoding HA–I53_dn5B (1000 µg per 1 L of cells). Expifectamine was used as transfection reagent according to the manufacturer's protocol. Cells were incubated for 96 hours, after which they were spun down by centrifugation (4,000 RPM, 20 min, 4 °C) and the protein-containing supernatant was further clarified by vacuum-filtration (0.45 µm, Millipore Sigma). For nickel-affinity chromatography steps, a background of 50 mM Tris, 350 mM NaCl, pH 8.0 was added to clarified supernatant. For each liter of supernatant, 4 mL of Ni Sepharose excel resin (GE) was rinsed into phosphate-buffered saline (PBS) using a gravity column and then added to the supernatant, followed by overnight shaking at 4 °C. After 16-24 hours, resin was collected and separated from the mixture and washed twice with 50 mM Tris, 500 mM NaCl, 30 mM imidazole, pH 8.0 prior to elution of desired protein with 50 mM Tris, 500 mM NaCl, 300 mM imidazole, pH 8.0. Eluates were concentrated and applied to a HiLoad 16/600 Superdex 200 pg column pre-equilibrated with PBS for gel filtration.

**Production and Purification of DS-Cav1–Fusion Nanoparticles.** Gene synthesis and cloning was performed by Genscript. The protein sequence for DS-Cav1–I53_dn5b is shown in Supplementary Table 8. HEK 293F cells were grown in suspension using Expi293 expression medium (Thermo Fisher Scientific) at 150 RPM, 8% $CO_2$, 70% humidity, 37 °C. At confluency of ~2.5 to $3\times10^6$ cells/ml, the cells were transiently transfected with the vector encoding DS-Cav1–I53_dn5B (1 mg per 1 L of cells). Expifectamine was used as transfection reagent according to the manufacturer's protocol. Cells were incubated for 96 hours and spun down by centrifugation (4,000 RPM for 20 minutes at 4 °C). Supernatant was vacuum-filtered (0.45 µm, Millipore Sigma) and 50 mM Tris, 350 mM NaCl, pH 8.0 was added for nickel-affinity chromatography. Ni Sepharose resin (GE) was washed three times with PBS by centrifugation (2,000 RPM for 5 minutes at 4 °C) and added to the supernatant. Nickel-supernatant was incubated either overnight at 4 °C or for 2 hours at room temperature. Resin was collected and separated from the mixture and washed twice with 50 mM Tris, 500 mM NaCl, 30 mM imidazole, pH 8.0 prior to elution of desired protein with 50 mM Tris, 500 mM NaCl, 300 mM imidazole, pH 8.0. Eluates were concentrated and applied to a HiLoad 10/300 Superdex 200 Increase GL column pre-equilibrated with PBS size exclusion chromatography.

**Assembly and Purification of Antigen-Fused Nanoparticles.** Several reactions containing 5-10 µg of the purified fusion component and an equimolar amount of the counter bare assembly component were prepared and incubated under different conditions (varying temperature and assembly buffer) for 24 hours. Native PAGE Bis-Tris gels (Thermo Fisher Scientific) and negative stain electron microscopy was used for detection of assembly efficacy. Following the identification of optimal assembly conditions, milligram quantities of particles were assembled and purified using gel filtration chromatography (Superose 6 or Sepharose 500 column) with TBS as the running buffer. Fractions corresponding to the fusion component were pooled and concentrated (Amicon Ultra Centrifugal Filter Units, Millipore Sigma).

**Biolayer Interferometry on HA–I53_dn5.** To produce biotin-labeled antibodies specific to the H1 HA head, 5J8 monoclonal antibody (mAb)[18] in PBS was mixed with a 20× molar excess (relative to complete antibodies) of EZ-Link™ NHS-LC-Biotin (Thermo Fisher Scientific) and allowed to sit for 2 hours at 4 °C, followed by two rounds of overnight dialysis against PBS at 4 °C to remove excess biotinylation reagent. All biosensors were hydrated in assay buffer (25 mM Tris, 150 mM NaCl, 0.5% bovine serum albumin, 0.01% TWEEN 20, pH 8.0) before use. Biotinylated 5J8 (0.02 mg/mL in assay buffer) was immobilized on SA biosensors, then briefly dipped in assay buffer prior to exposure to designed H1 HA fusions (500 nM per asymmetric unit, in assay buffer). The biosensor was again dipped in assay buffer and then exposed to the stem-specific CR6261 mAb[19].

**ELISA Assays on DS-Cav1–I53_dn5.** ELISA was used to measure binding kinetics of DS-Cav1–I53_dn5 to RSV F-specific mAbs D25, Motavizumab, and AM14. D25 is a pre-fusion specific mAb that binds site Ø[20]. Motavizumab binds site II of the pre and post-fusion conformations. AM14 is trimer-specific binding across protomers of pre-F. 96-well enzyme-linked immunosorbent assay (ELISA) plates were coated with 2 µg/mL DS-

Cav1/H1 HA nanoparticles. Plates were incubated at 4 °C overnight and blocked with PBS containing 5% skim milk at 37 °C for 30 minutes. mAbs listed above serially diluted in fourfold steps, were then added and the plates incubated at 37 °C for 45 minutes. Horseradish peroxidase (HRP)-conjugated anti-human IgG (Southern Biotech., Birmingham, AL) was added and incubated at 37 °C for 30 minutes, followed by 3,3′,5′,5-Tetramethylbenzidine (TMB; Sigma-Aldrich, St. Louis, MO) HRP substrate, and yellow color that developed after the addition of 1 M $H_2SO_4$ was measured by absorbance at 450 nm.

### Supplementary Figures

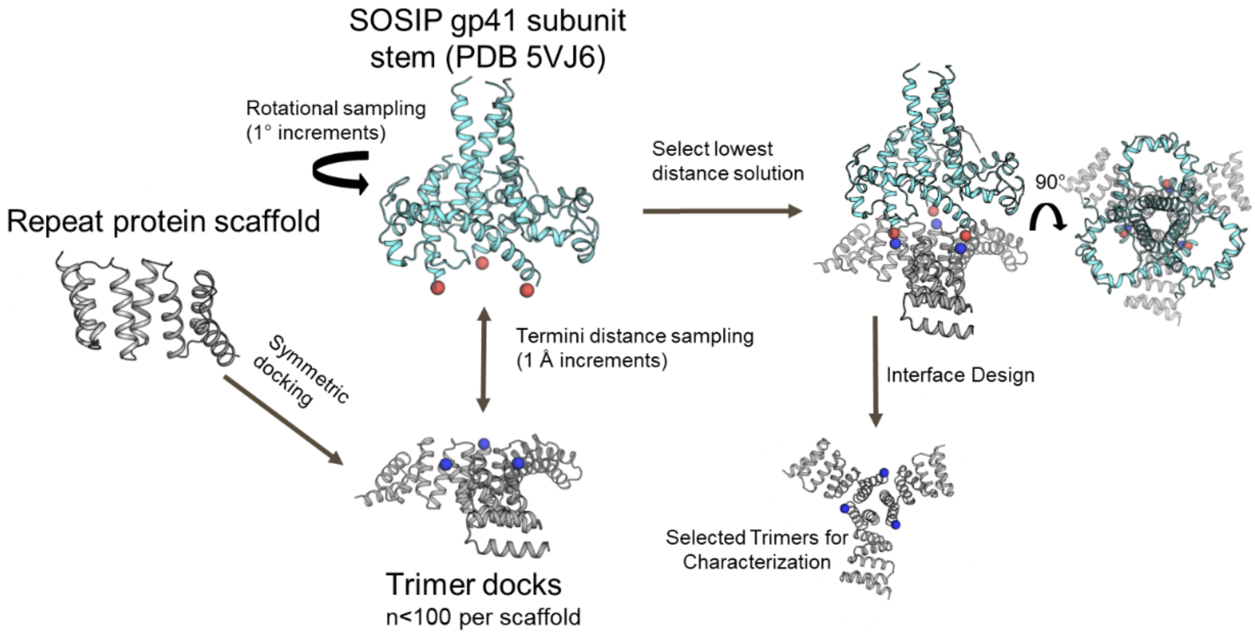

**Supplementary Figure 1.** *In silico* trimer docking and antigen targeting strategy. Designs for C3-symmetric trimers (N terminal residue labeled in blue) capable of scaffolding target antigens shown in gray. BG505 SOSIP (C terminal residue 664 labeled in red) trimeric glycoprotein subunit gp41 shown in cyan for illustration of targeting strategy.

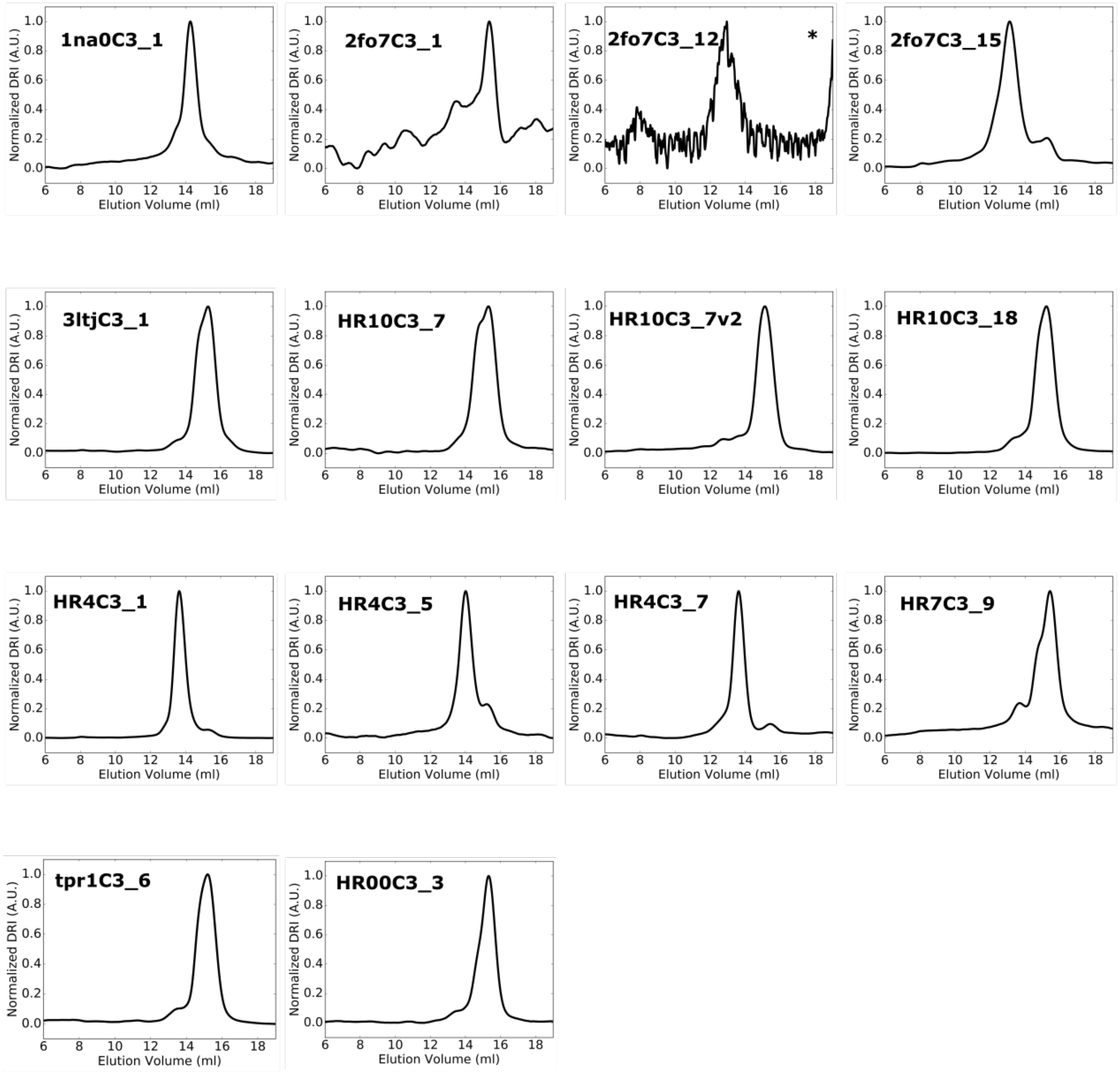

*Chromatogram obtained on Superdex 75 10/300 GL gel filtration column.

**Supplementary Figure 2.** SEC-MALS chromatograms for failed designs, occupying an unintended oligomeric configuration. Predominant oligomeric species for each design were collected by fractionation from a primary size exclusion run, and 14 sizing profiles are presented here from the subsequent round of high-performance size exclusion chromatography from a Superdex 200 gel filtration column.

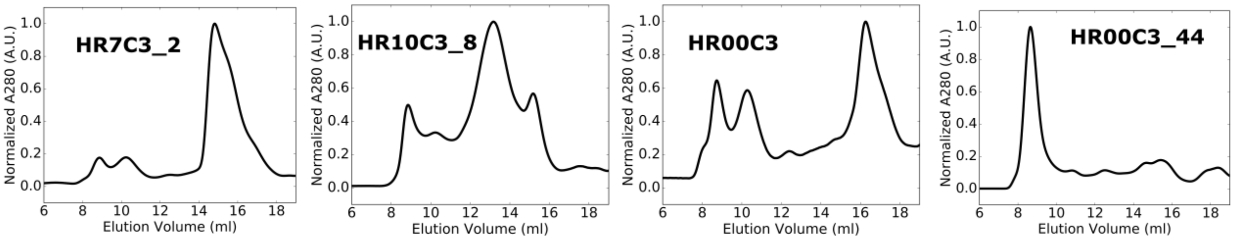

**Supplementary Figure 3.** SEC chromatograms for trimers with off-target retention volumes after $Ni^{2+}$ IMAC. Primary size exclusion chromatograms obtained from a Superdex 200 gel filtration column for soluble proteins directly after purification by $Ni^{2+}$ IMAC. 2 designs displayed polydisperse profiles indicating formation of off-target oligomers, whereas 4 designs formed soluble aggregate.

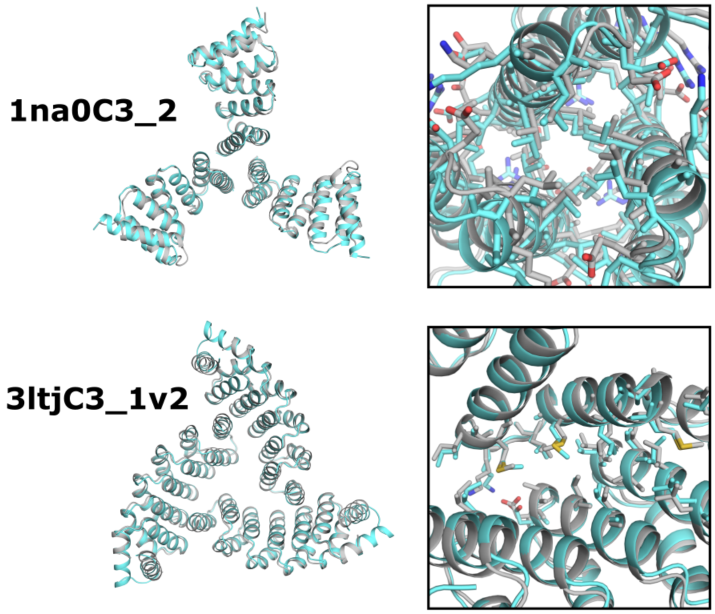

**Supplementary Figure 4.** Comparison between the experimentally determined crystal structures and corresponding models of designed trimers. A full model and crystal structure superposition is displayed, with crystal structures shown in cyan and models in gray. Magnified view illustrates the side chains at the designed interface.

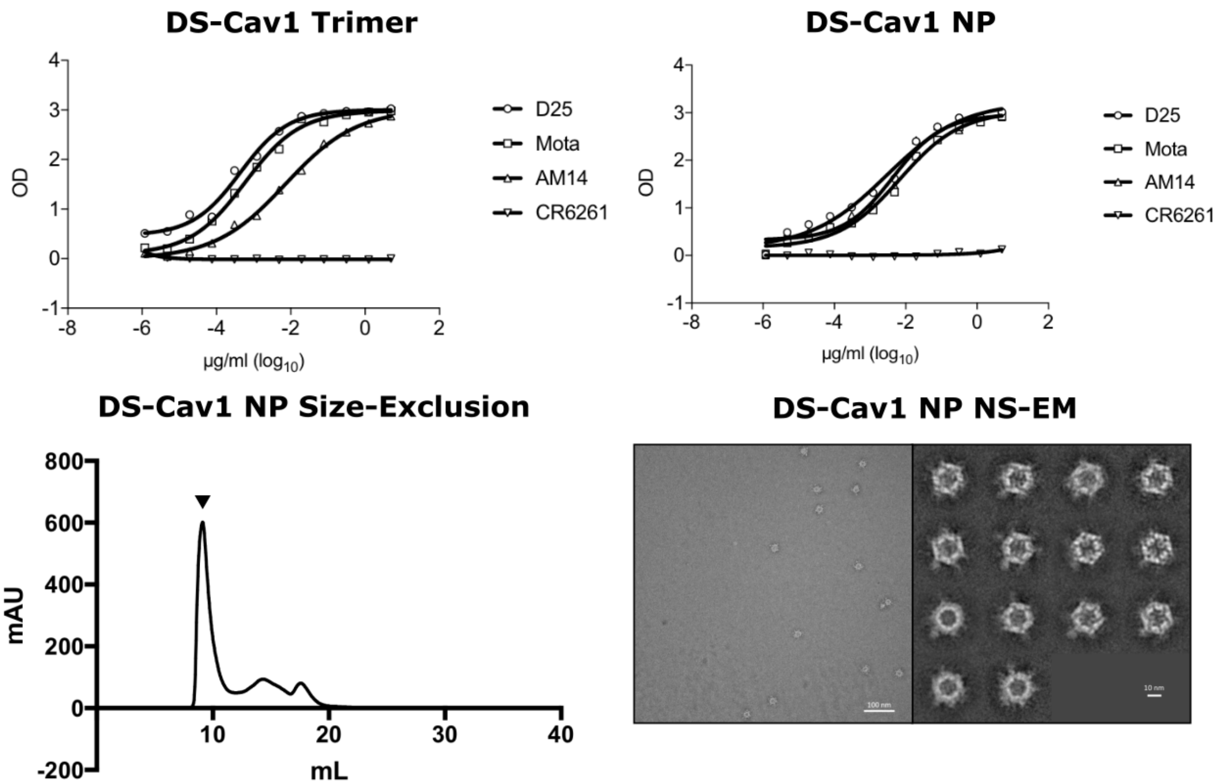

**Supplementary Figure 5.** Top panel: ELISA using anti-DS-Cav1 antibodies bound to DS-Cav1 trimer with foldon or designed nanoparticle I53_dn5. Fluorescence signal is plotted as a function of binding to purified trimer and nanoparticle. Bottom panel: size-exclusion profile of DS-Cav1–I53_dn5 particle assembly on a Superose 6 column and negative stain electron micrograph field image and 2D class averages of purified particle.

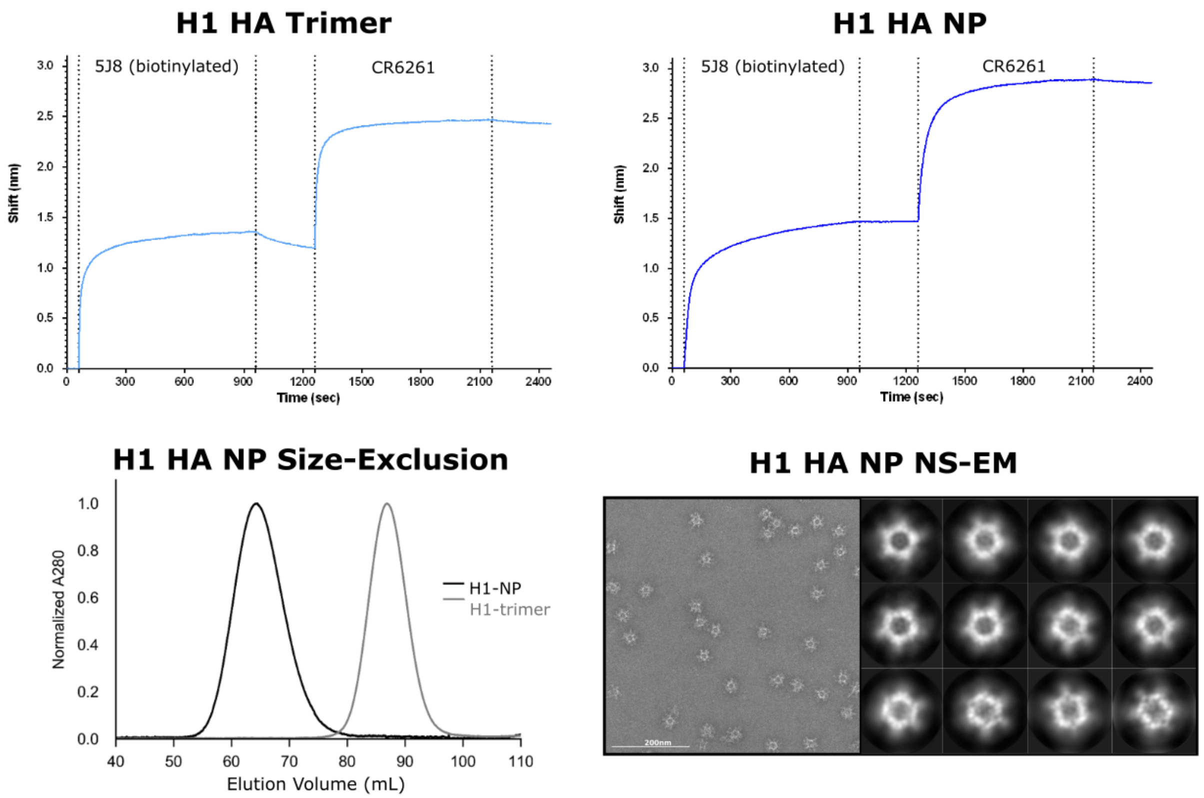

**Supplementary Figure 6.** Top panel: Octet bio-layer interferometry using plate-coated head-directed H1 HA mAb (5J8) for antigen capture, and subsequent stem-directed H1 HA mAb (CR6261) addition to both HA–I53_dn5 (NP) and corresponding trimeric component HA–I53_dn5B (trimer). Bottom panel: size-exclusion profile of HA–I53_dn5 assembled particle and corresponding trimer HA–dn5B assembly on a Sephacryl S-500 column, and negative stain electron micrograph field image and 2D class averages of purified particle.

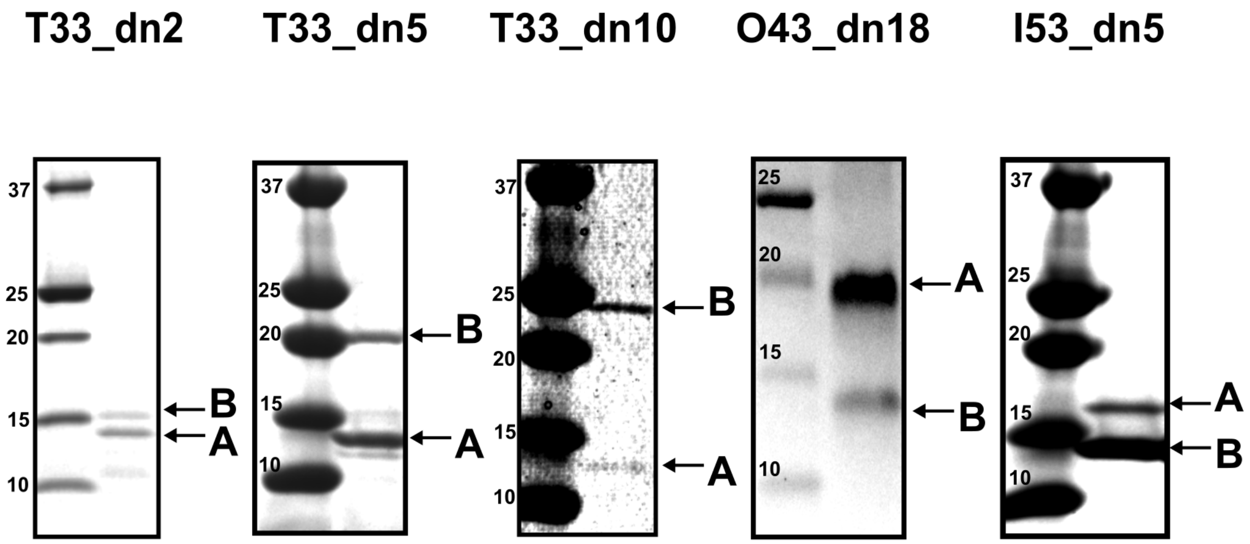

**Supplementary Figure 7.** SDS-PAGE of bicistronically-expressed de novo nanoparticle eluted from $Ni^{2+}$ IMAC. For each designed nanoparticle: Left - protein standard (Precision Plus Dual Xtra, Bio-Rad). Right - labeled bands corresponding to the expected size of each component.

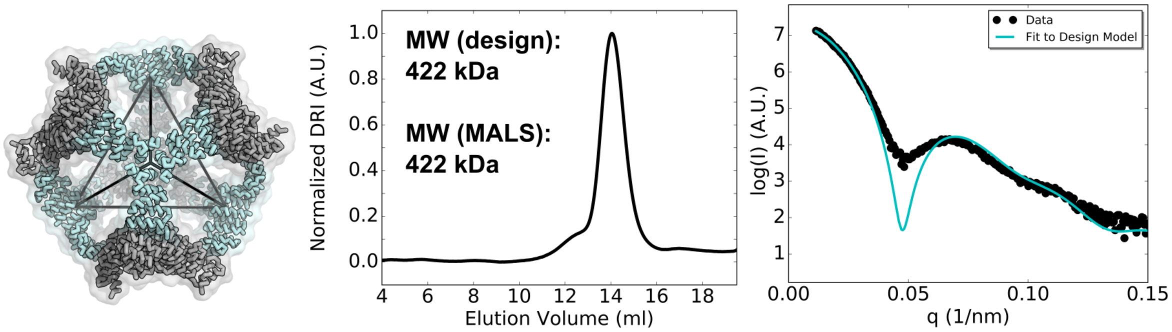

**Supplementary Figure 8.** Biophysical Characterization of T33_dn5. Left - designed model. Middle - size-exclusion chromatograms and calculated molecular weights from multi-angle light scattering. Right - small-angle X-ray scattering comparisons between experimental data and profile computed from model.

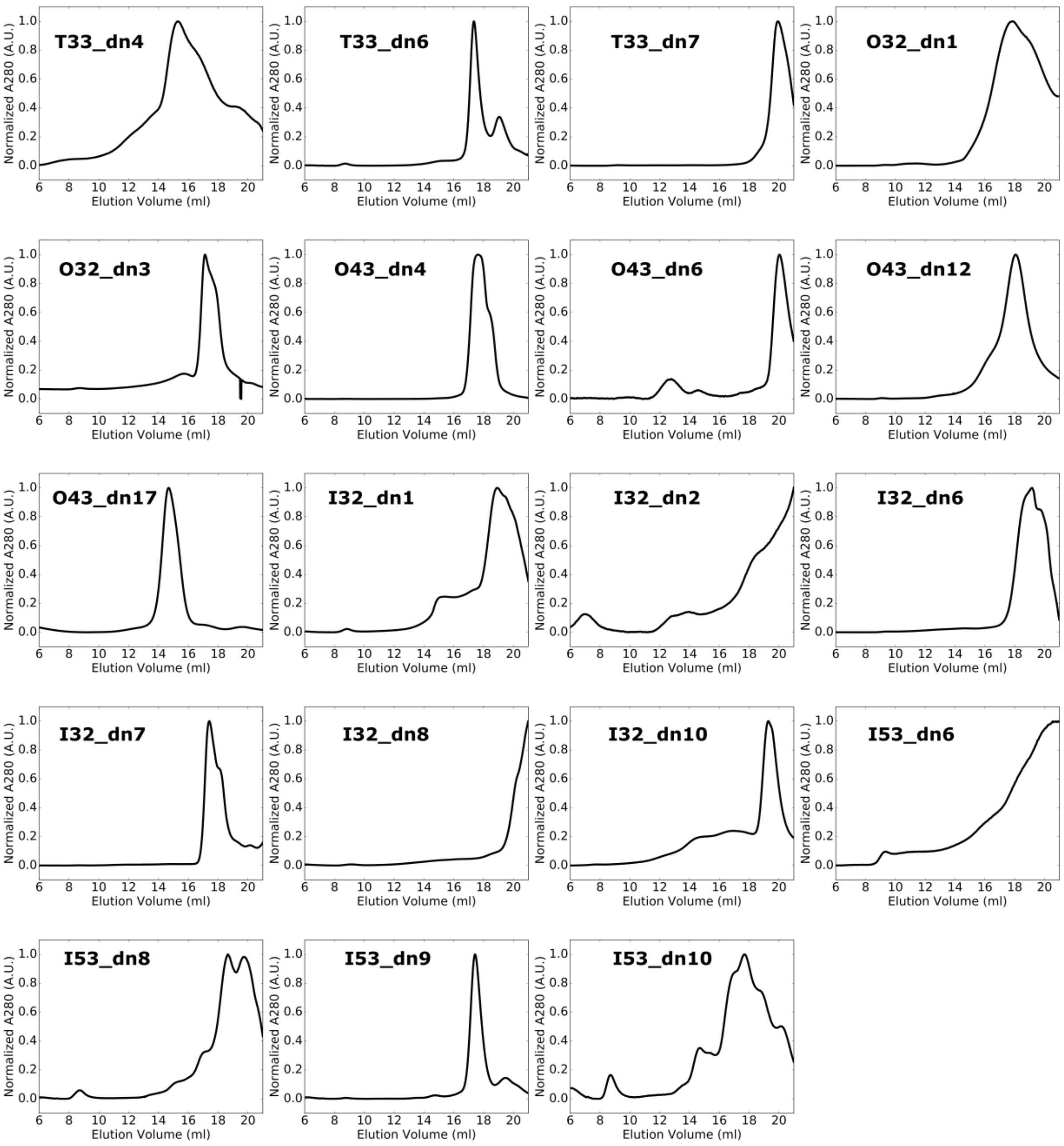

**Supplementary Figure 9.** SEC profiles for two-component nanoparticles that failed to co-assemble as predicted by Rosetta. Primary size exclusion chromatograms obtained from a Superose 6 gel filtration column for soluble proteins directly after purification by $Ni^{2+}$ IMAC. Designs presented here appear to form off-target assemblies based on retention volume or occupy polydisperse complex states.

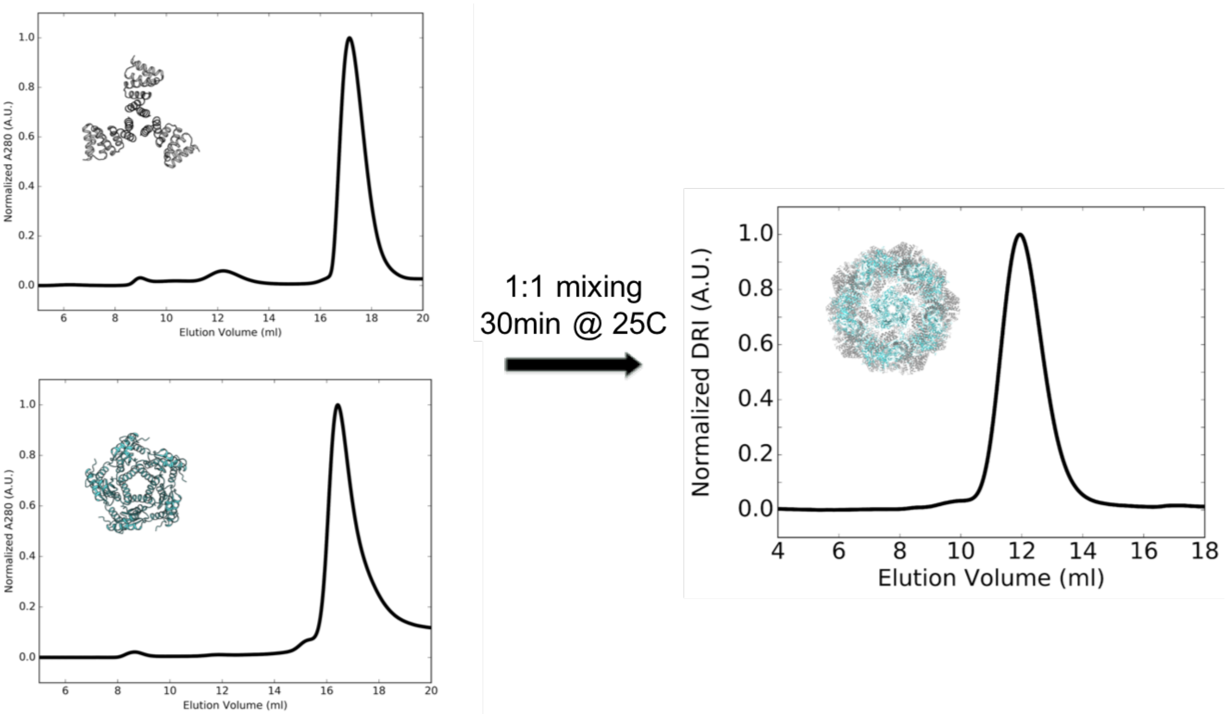

**Supplementary Figure 10.** Size-exclusion chromatogram of individual components I53_dn5A (pentamer) and I53_dn5B (trimer) and assembly run on a Superose 6 10/300 GL column.

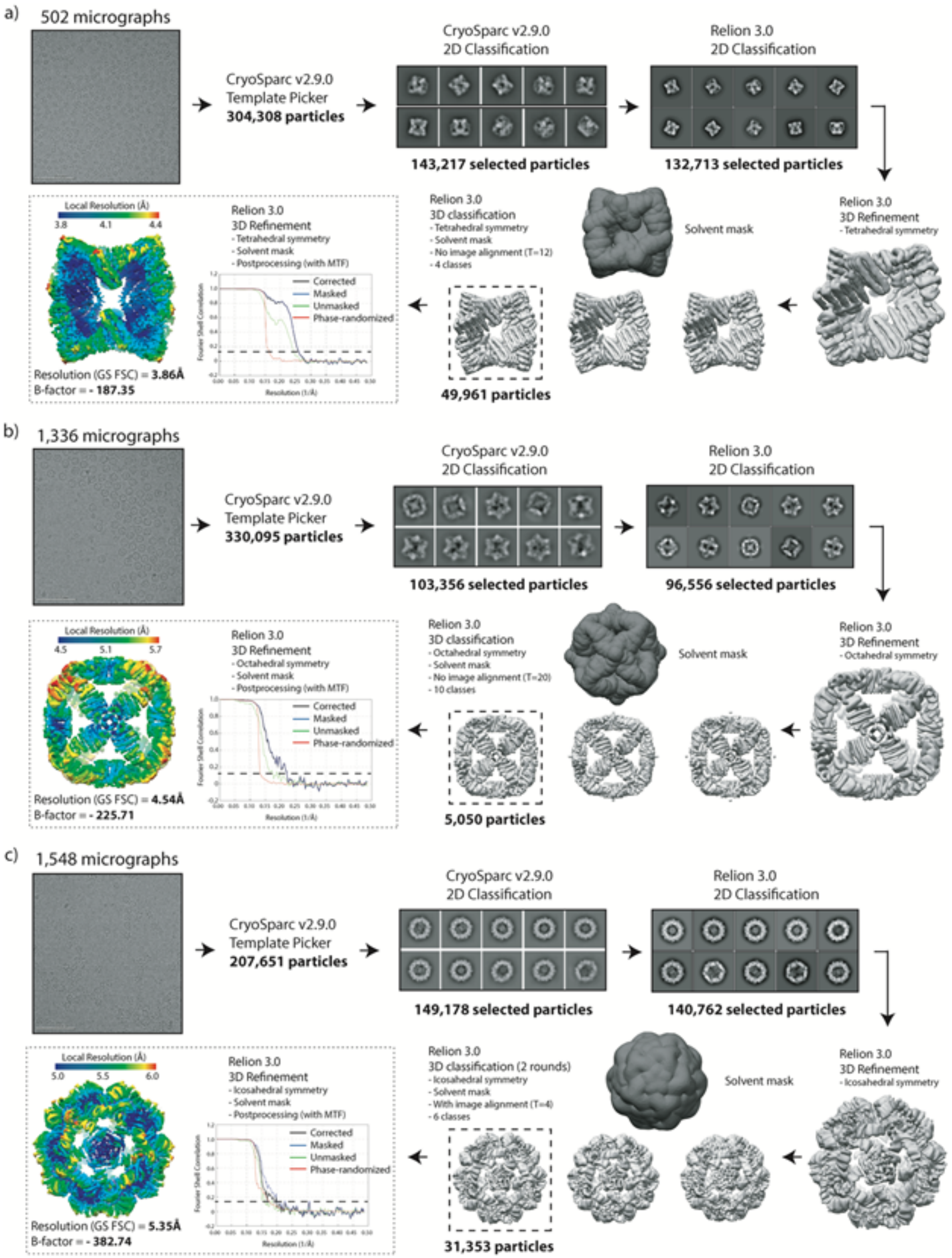

**Supplementary Figure 11.** Cryo-electron microscopy data processing workflow and relevant statistics for a) T33_dn10, b) O43_dn18, and c) I53_dn5.

**Supplementary Tables**

| Design | Targeted Antigens | Experimental Molecular Weight (kDa) | Target Molecular Weight (kDa) | SAXS $X$ value | Resolution, r.m.s.d. structure (Å, Å) |
|---|---|---|---|---|---|
| 1na0C3_2 | HA, SOSIP, DS-Cav1 | 48 | 45 | 1.4 | 2.6, 1.4 |
| 3ltjC3_1v2 | SOSIP, DS-Cav1 | 56 | 63 | 1.1 | 2.3, 0.8 |
| 3ltjC3_11 | SOSIP, DS-Cav1 | 50 | 66 | 1.6 | -- |
| HR04C3_5v2 | SOSIP | 71 | 69 | 1.5 | -- |
| T33_dn2 | HA, SOSIP, DS-Cav1 | 397 | 345 | 4.8 | -- |
| T33_dn5 | HA, SOSIP, DS-Cav1 | 422 | 422 | 1.7 | -- |
| T33_dn10 | HA, SOSIP, DS-Cav1 | 546 | 556 | 2.3 | 3.9, 0.65 |
| O43_dn18 | HA,SOSIP, DS-Cav1 | 810 | 876 | 2.9 | 4.5, 0.98 |
| I53_dn5 | HA, SOSIP, DS-Cav1 | 2000 | 1960 | 1.2 | 5.3, 1.30 |

**Supplementary Table 1.** Summary of the experimental results from characterization of the antigen-tailored symmetric homotrimeric proteins and subsequent two-component nanoparticles. 1na0C3_2 and 3ltjC3_1v2 structures determined by X-ray crystallography and T33_dn10, O43_dn18, and I53_dn5 determined by cryo-electron microscopy.

| Design | Target Molecular Weight (kDa) | Experimental Molecular Weight (kDa) | Approximate Oligomerization State |
|---|---|---|---|
| 1na0C3_1 | 44 | 43 | 3 |
| 2fo7C3_1 | 51 | 20 | 1 |
| 2fo7C3_12 | 50 | 106 | 6 |
| 2fo7C3_15 | 51 | 70 | 4 |
| 3ltjC3_1 | 63 | 37 | 2 |
| HR10C3_7 | 67 | 47 | 2 |
| HR10C3_7v2 | 67 | 38 | 2 |
| HR10C3_18 | 67 | 35 | 2 |
| HR4C3_1 | 69 | 75 | 3 |
| HR4C3_5 | 69 | 61 | 3 |
| HR4C3_7 | 68 | 65 | 3 |
| HR7C3_9 | 56 | 30 | 2 |
| tpr1C3_6 | 48 | 28 | 2 |
| HR00C3_3 | 94 | 37 | 1 |

**Supplementary Table 2.** SEC-MALS data for designs intended to be homotrimeric with C3 symmetry.

| Design | MW A (kDa) | MW B (kDa) | MW model (kDa) | MW exp (kDa) | Rg model (Å) | Rg exp (Å) | Dmax model (Å) | Dmax exp (Å) | *X* | qmax (1/nm) |
|---|---|---|---|---|---|---|---|---|---|---|
| **1na0C3_2** | 14.99 | - | 44.97 | 48 | 26.4 | 29.5 | 84 | 86 | 1.4 | 0.23 |
| **3ltjC3_1v2** | 20.90 | - | 62.69 | 56 | 27.7 | 31.3 | 88 | 94 | 1.1 | 0.18 |
| **3ltjC3_11** | 22.21 | - | 66.62 | 50 | 28.3 | 30.3 | 87 | 92 | 1.6 | 0.20 |
| **HR04C3_5v2** | 23.05 | - | 69.14 | 71 | 25.9 | 28.6 | 82 | 86 | 1.5 | 0.25 |
| **T33_dn2** | 13.82 | 14.88 | 344.45 | 397 | 61.4 | 64.7 | 169 | 169 | 4.8 | 0.17 |
| **T33_dn5** | 13.72 | 21.49 | 422.42 | 422 | 66.7 | 69.4 | 177 | 193 | 1.7 | 0.16 |
| **T33_dn10** | 14.07 | 31.42 | 545.88 | 556 | 62.3 | 60.1 | 169 | 170 | 2.3 | 0.20 |
| **O43_dn18** | 22.69 | 13.82 | 876.26 | 810 | 80.6 | 81.3 | 217 | 221 | 2.9 | 0.28 |
| **I53_dn5** | 17.19 | 15.33 | 1951.57 | 2000 | 95.9 | 97.1 | 241 | 243 | 1.2 | 0.21 |

**Supplementary Table 3.** Biophysical properties of designed trimers (top) and two-component nanoparticles (bottom). Experimentally-measured data (exp) is compared to predicted design data (model). Molecular weights (MW) were obtained using the ASTRA software. $R_g$ and $D_{max}$ calculations performed in Scatter3 SAXS analysis software with the determined $q_{max}$ values. *X* values computed from the FoXS online SAXS web server between the designed model and the experimental scattering data.

| | 1na0C3_2 (PDB ID X) | 3ltjC3_1v2 (PDB ID Y) |
|---|---|---|
| **Data Acquisition** | | |
| Space group | P 1 21 1 | R 3 :H |
| Cell dimensions<br>a, b, c (Å)<br>α , β , γ (°) | 69.47, 64.91, 99.03<br>90, 106.33, 90 | 88.182, 88.182, 65.244<br>90, 90, 120 |
| $R_{merge}$ | 0.1427 (0.8023) | 0.1189 (1.041) |
| $CC_{1/2}$ | 0.994 | 0.997 (0.623) |
| $\langle I/\sigma_I \rangle$ | 9.5 (2.16) | 9.35 (1.30) |
| Completeness (%) | 99.86 | 91.51 (77.88) |
| Multiplicity | 4.6 (4.6) | 5.6 (5.1) |
| Wilson B-factor (Å$^2$) | 37.11 | 40.43 |
| **Refinement** | | |
| Resolution range (Å) | 38.87 - 2.53 (2.60 - 2.53) | 44.09 - 2.303 (2.386 - 2.303) |
| No. of reflections for refinement | 27066 (2030) | 8361 (655) |
| No. of reflections for $R_{free}$ | 1442 (103) | 756 (62) |
| $R_{work}$(%)/$R_{free}$(%) | 0.1859 (0.195) / 0.2316 (0.240) | 0.2021 (0.2555) / 0.2261 (0.2240) |
| Water count | 110 | 25 |
| Residue count | 711 | 180 |

| | | |
|---|---|---|
| Average B-factors (Å$^2$)<br>Protein<br>Water | <br>47.40<br>46.01 | <br>49.36<br>50.80 |
| r.m.s.d. deviations<br>Bond length (Å)<br>Bond angles (°) | <br>0.68<br>0.78 | <br>0.002<br>0.44 |
| Ramachandran favored (%) | 98.00 | 100.00 |
| Ramachandran allowed (%) | 1.00 | 0.00 |
| Ramachandran outliers (%) | 1.00 | 0.00 |
| Rotamer outliers (%) | 3.00 | 0.00 |

**Supplementary Table 4.** Crystallography data collection and refinement statistics for constructs 1na0C3_2 and 3ltjC3_1v2. Statistics for the highest-resolution shell are shown in parentheses.

| | T33_dn10 | O43_dn18 | I53_dn5 |
|---|---|---|---|
| Microscope | Titan Krios | Titan Krios | Titan Krios |
| Voltage (kV) | 300 | 300 | 300 |
| Detector | Gatan K2 Summit | Gatan K2 Summit | Gatan K2 Summit |
| Recording mode | Counting | Counting | Counting |
| Magnification | 29,000 X | 29,000 X | 29,000 X |
| Movie micrograph pixel size | 1.03 | 1.03 | 1.03 |
| Dose rate ($e^-/Å^2/s$) | 5.04 | 5.04 | 4.46 |
| No. of frames per movie micrograph | 40 | 40 | 45 |
| Frame exposure time (ms) | 250 | 250 | 250 |
| Movie micrograph exposure time (s) | 10.00 | 10.00 | 11.25 |
| Total dose ($e^-/Å^2$) | 50.4 | 50.4 | 50.2 |
| Under focus range (μm) | 0.6 - 1.6 | 0.6 - 1.6 | 0.6 - 1.6 |
| Number of movie micrographs | 502 | 1,336 | 1,548 |

**Supplementary Table 5.** Cryo-electron microscopy data acquisition metrics for nanoparticle constructs T33_dn10, O43_dn18, and I53_dn5.

| | T33_dn10 | O43_dn18 | I53_dn5 |
|---|---|---|---|
| PDB | 6VFH | 6VFI | 6VFJ |
| Residues | 4,752 | 7,560 | 16,320 |
| Amino-acids | 4,752 | 7,560 | 16,320 |
| Carbohydrates | 0 | 0 | 0 |
| RMSD Bonds | 0.019 | 0.018 | 0.020 |
| RMSD Angles | 1.389 | 1.484 | 1.648 |
| Ramachandran | | | |
| Favored (%) | 99.13 | 98.71 | 98.88 |
| Allowed (%) | 0.87 | 1.29 | 1.12 |
| Outliers (%) | 0.00 | 0.00 | 0.00 |
| Rotamer outliers | 0.00 | 0.00 | 0.00 |
| Clash score | 0.32 | 0.41 | 0.27 |
| Molprobity score | 0.62 | 0.65 | 0.60 |
| EMRinger score | 2.12 | 0.68 | 0.69 |

**Supplementary Table 6.** Cryo-electron microscopy model building and refinement statistics for nanoparticle designs T33_dn10, O43_dn18, and I53_dn5.

| Design | Sequence |
|---|---|
| **1na0C3_1<br>(SEC-MALS)** | **MNIAEAAYRVGNKAYKKGRYELAILAYILAILLDPNNAEAWYNLGNAYYKEG<br>EYDEAIEYYQKALELDPNNAEAWYNLGNAYYKQGDYDEAIEYYQKALELDP<br>NNAEAKQNLGNAKQKQGLEHHHHHH** |
| **1na0C3_2<br>(SEC-MALS, SAXS)** | **MEEAELAYLLGELAYKLGEYRIAIRAYRIALKRDPNNAEAWYNLGNAYYKQG<br>DYDEAIEYYQKALELDPNNAEAWYNLGNAYYKQGDYDEAIEYYQKALELDP<br>NNAEAKQNLGNAKQKQGLEHHHHHH** |
| **2fo7C3_1<br>(SEC-MALS)** | **MAERLYKLGNKAYKRGEYILALIAYVVALRDDPRSAEAWYNLGNAAYKSGE<br>YDEAIEYYQKALELDPRSAEAWYNLGNAYYKQGDYDEAIEYYQKALELDPR<br>SAEAWYNLGNAYYKQGDYDEAIEYYQKALELDPRSLEHHHHHH** |
| **2fo7C3_12<br>(SEC-MALS)** | **MAEKAYNIGNAAYKEGEYRVAILAYMLALLADPRSAEALYNLGNAAYKEGD<br>YKVAIAAYLLALDLDPRSAEAWYNLGNAYYKQGDYDEAIEYYQKALELDPR<br>SAEAWYNLGNAYYKQGDYDEAIEYYQKALELDPRSLEHHHHHH** |
| **2fo7C3_15<br>(SEC-MALS)** | **MALRWLLLGILAMLLGAEELAIEAYQKALELEPRSAMAWLALGAAYYKEGD<br>YDEAIEYYQKALELDPRSAAAWALLGNAYYKQGDYDEAIEYYQKALENRPR<br>SARAWYNLGNAYYKQGDYDEAIEYYQKALELDPRSLEHHHHHH** |
| **3ltjC3_1<br>(SEC-MALS)** | **MTDPLAVILYIAILKAEKSIARAKAAEALGKIGDERAVEPLIKALKDEDALVRA<br>AAADALGQIGDERAVEPLIKALKDEEGLVRASAAIALGQIGDERAVEPLIKAL<br>KDERDLVRVAAAVALGRIGDERAVEPLIKALKDEEGEVREAAAIALGSIGGE<br>RVRAAMEKLAETGTGFARKVAVNYLETHKLEHHHHHH** |
| **3ltjC3_1v2<br>(SEC-MALS, SAXS)** | **MTDPMKVILYIAMLELEKYIMRAAAAYALGKIGDERAVEPLIKALKDEDAIVR<br>AAAADALGQIGDERAVEPLIKALKDEDGAVRVSAAVALGQIGDERAVEPLIK<br>ALKDEDAVVRVAAAIALGLIGDERAVEPLIKALKDEKGKVREAAALALGAIG<br>GERVRAAMEKLAETGTGFARKVAVNYLETHKLEHHHHHH** |
| **3ltjC3_11<br>(SEC-MALS, SAXS)** | **MRREETDPLAVVMYRLNLRDDSYYVRRAAAYALGKIGDERAVEPLIKALKD<br>EDAWVRRAAADALGQIGDERAVEPLIKALKDEDGWVRQSAAVALGQIGDE<br>RAVEPLIKALKDEDWFVRAAAAAALGRIGDERAVEPLIKALKDEDEMVREIA<br>ALALGMIGGERVRAAMEKLAETGTGFARKVAVNYLETHKLEHHHHHH** |
| **HR4C3_1<br>(SEC-MALS)** | **MDICELEARLVALLVLLAKRAGADEDLIAELVAVMIMIVILRLKKSGSSYEVIC<br>ECVARIVAEIVEALKRSGTSEDEIAEIVARVISEVIRALKRSGSSYEVICECVAR<br>IVAEIVEALKRSGTSEDEIAEIVARVISEVIRTLKESGSSYEVIKECVQRIVEEIV<br>EALKRSGTSEDEINEIVRRVKSEVERTLKESGSLEHHHHHH** |
| **HR4C3_5<br>(SEC-MALS)** | **MDECEEKARRVAEKVERLKRSGTSEDEIAEEVAREISEVIRTLKESGSEYKVI<br>CRCVARIVAEIVEALKRSGTSEDEIAEIVARVISEVIRTLKESGSKYKIICICVAII<br>VAEIVAALKRSGTSEDEIAEIVARVISEVIRTLKESGSSYEVIKQCVQAIVQAIIL<br>ALMKSGTEVEEILLIVLRVEEEVERTLKESGSLEHHHHHH** |
| **HR4C3_5v2<br>(SEC-MALS, SAXS)** | **MDECEEKARRVAEKVERLKRSGTSEDEIAEEVAREISEVIRTLKESGSEYKVI<br>CRCVARIVAEIVEALKRSGTSEDEIAEIVARVISEVIRTLKESGSDYLIICVCVAII<br>VAEIVEALKRSGTSEDEIAEIVARVISEVIRTLKESGSSYEVIKECVQIIVLAIILA<br>LMKSGTEVEEILLILLRVKTEVRRTLKESGSLEHHHHHH** |
| **HR4C3_7<br>(SEC-MALS)** | **MDECEKKARLVAILVIVAKALGAEEKLIALLVALEIVVVIIELKASGSSYEVICE<br>CVARIVAEIVEALKRSGTSEDEIAEIVAKVIAAVIIVLKELGSSYEVICECVARIV** |

| | |
|---|---|
| | **AEIVEALKRSGTSEDEIAEIVARVISEVIRTLKESGSSYEVIKECVQRIVEEIVEALKRSGTSEDEINEIVRRVKSEVERTLKESGSLEHHHHHH** |
| **HR7C3_2 (SEC)** | **MEKRIARELCELAAERAAESNDEREARIAAIECLLVAERAGMPTKEAARSFCEAAARAAAESNDEEVAKIAAKACLEVAKQAGMPTKEAARSFCEAAARAAAESNDEEVAKIAAKACLEVAKQAGMPTKEAARSFCEAAKRAAKESNDEEVEKIAKKACKEVAKQAGMPLEHHHHHH** |
| HR7C3_8 | MTEEDAARTCKKAARKAAESNDEEVAKQAAKDCLEVAKQAGMPTTIAAAIFCLAAARAAAESNDEEVAKIAAKACLEVAKQAGMPTKAAAIAFCIAAAMAAAESRDEEVAKIAAKACLEVAKQAGMPTKTAAALFMIAAIAAALRSEDEVVLAIAALAIAEVLKQAGMPLEHHHHHH |
| **HR7C3_9 (SEC-MALS*update)** | **MTKEMAAVLCMVLALKAAESNDEEKAKKAAKLCLIMADEAGMPTKEAARSFCEAAAIAAAVSEDEEVAKIAAKACLEVAKQAGMPTKEAARSFCEAAASAAAILNEEEVAKIAAKACLEVAKQAGMPTKEAARSFCEAAKRAAKRSNDEEVEKIAKKACKEVAKQAGMPLEHHHHHH** |
| **HR10C3_7 (SEC-MALS)** | **MSSEKEELRELLVAIVAVAAEDKGDDTEEAREAAREAFELVREAAERAGIDSSEVLTLAILLILIVVLIAADAGYDISEAARAAAEAFKRVAEAAKRAGITSSEVLELAIRLIKEVVVNAAIRGYDISEAARAAAEAFKRVAEAAKRAGITSSKILKMAIILIRVMVKMAKERGKDISEAARQAAEIFRKAAERMRGSLEHHHHHH** |
| **HR10C3_7v2 (SEC-MALS)** | **MSSEKEELRKMLVALVVVAAKEKGDDTEEAREAAREAFELVREAAERAGIDSSVVLALAILLILLVVLAAQMAGYDISEAARAAAEAFKRVAEAAKRAGITSSEVLELAIRLIKRVVLNAQIRGYDISEAARAAAEAFKRVAEAAKRAGITSSLLLKMAIVLIRVLVELAQESGADISEAARKAAEIMRRAAEDMRGSLEHHHHHH** |
| **HR10C3_8 (SEC)** | **MDECEEKARRVAEKVERLKRSGTSEDEIAEEVAREISEVIRTLKESGSEEVEICACVARIVAEIVEALKRSGTSEDEIAEIVARVISEVIRTLKESGSSYLVICMCVALIVAQIVEALKRSGTSRKEIAEIVARVISEVIRTLKESGSSYEVIKECVERIVRAIVLALRESGTRITEIMAIVLAVLKEVLRTLKESGSLEHHHHHH** |
| **HR10C3_18 (SEC-MALS)** | **MKREKMELAKRLLKIVVENAKRKGDEEALAALAALLAFALVREAAERAGIDSSEVLELAIRLIKEVVENAQREGYRIALAALVAAMAFAVVAEAAKEAGITSSEVLELAIRLIKEVVENAQREGYEIVDAAMAAALAFARVAEAAKRAGITSSETLKRAIEEIRKRVEEAQREGNDISEAAEQAAEEFRKKAEELKLEHHHHHH** |
| **HR00C3 (SEC)** | **MKEEKIAKLISLLAELSKKLIEIVARAADNKTTEEAVDIAILLIAIIARLAIRLIEMLAKNLASEEFMARAISAIAELAKKAIEAIYRLADNHTTDIRMLKAILAIAELAAEAIKAIADLAKNHTTEEFMARAISAIAELAKKAIEAIYRLADNHTTDLFMAIAIMAIAVLALLAIMAIADLAKNHTTEEFMAKAISAIAELAKKAIEAIYRLADNHTSPDLIELAILAIEIIALAAIIAIEELAENITTEEYKEKAKSAIDEIREKAKEAIKRLEDNRTLEHHHHHH** |
| **HR00C3_3 (SEC-MALS)** | **MKERLIAKLISVLAEASKILIRIAAKAADKLEREAAVILAIVLIAVAAIAAIAAIALLAANLASEEFMARAISAIAELAKKAIEAIYRLADNHTEDEAMALAIEIIAILALLAIVAIALLAANHTTEEFMARAISAIAELAKKAIEAIYRLADNHTTDTFMAKAIEAIAELAKEAIKAIAELAKNHTTEEFMAKAISAIAELAKKAIEAIYRLADNHTSPTYIEKAIEAIEKIARTAIKAIEDLAKNITTEEYKEKAKSAIDEIREKAKEAIKRLEDNRTLEHHHHHH** |
| **HR00C3_44 (SEC)** | **MTEEKIAKEISRIAEESKKRIEELARKADNKTREAVVALAIAKIALLAREAIKRIEDLAKNLASEEFMARAISAIAELAKKAIEAIYRLADNHTKDVLMLVAIVAIAELAKEAIKAIADLAKNHTTEEFMARAISAIAELAKKAIEAIYRLADNHRLVAAMLL** |

| | |
|---|---|
| | AIEAIAELAKEAIKAIADLAKNHTTEEFMAKAISAIAELAKKAIEAIYRLADNHR LPAAILLAALAILLIAVTAILAILILALNITTEEYKEKALSAIEEIVEKAEEAIDRLE DNLTLEHHHHHH |
| tpr1C3_6 (SEC-MALS) | MAEAWKELGKVLEKLGRLEEAAVAYLLAVIDDPNDAEAWKELGKVLEKLG ELDAAAVAYEAAIELDPNDAEAWKELGKVLEKLGRLRRAALAYIKAIALDPN DAEAWKELGKVAEKLGRLKIAARIYKKAIELDPNDLEHHHHHH |
| ank1C2_1* | MHHHHHHGSWGSSELGKRLIEAAENGNKDRVKDLIENGADVNASDSDGRT PLHHAAENGHAEVVALLIEKGADVNAKDSDGRTPLHHAAENGHDEVVLILLL KGADVNAKDSDGRTPLHHAAENGHKRVVLVLILAGADVNTSDSDGRTPLDL AREHGNEEVVKALEKQ |
| ank3C2_1* | MSELGKRLIEAAENGNKDRVKDLLENGADVNASDSDGKTPLHLAAENGHA KVVLLLLEQGADPNAKDSDGKTPLHLAAENGHAVVVALLLMHGADPNAKD SDGKTPLHLAAENGHEEVVILLLAMGADPNTSDSDGRTPLDLAREHGNEEV VKVLEDHGGWLEHHHHHH |
| 1na0C3_3* | MNLAEKMYKAGNAMYRKGQYTIAIIAYTLALLKDPNNAEAWYNLGNAAYKK GEYDEAIEAYQKALELDPNNAEAWYNLGNAYYKQGDYDEAIEYYQKALELD PNNAEAKQNLGNAKQKQGLEHHHHHH |
| 1na0C3_7* | MNSAEAMYKMGNAAYKQGDYILAIIAYLLALEKDPNNAEAWYNLGNAAYKQ GDYDEAIEYYQKALELDPNNAEAWYNLGNAYYKQGDYDEAIEYYQKALELD PNNAEAKQNLGNAKQKQGLEHHHHHH |
| HR00C3_2* | MIEEVVAEMIDILAESSKKSIEELARAADNKTTEKAVAEAIEEIARLATAAIQLI EALAKNLASEEFMARAISAIAELAKKAIEAIYRLADNHTTDTFMARAIAAIANL AVTAILAIAALASNHTTEEFMARAISAIAELAKKAIEAIYRLADNHTTDKFMAA AIEAIALLATLAILAIALLASNHTTEEFMAKAISAIAELAKKAIEAIYRLADNHTS PTYIEKAIEAIEKIARKAIKAIEMLAKNITTEEYKEKAKSAIDEIREKAKEAIKRL EDNRTLEHHHHHH |
| 1na0C4_1* | MTLARVAYILGAIAYAQGEYDIAITAYQVALDLDPNNAEAWYNLGNAYYKQG DYDEAIEYYQKALELDPNNAEAWYNLGNAYYKQGDYDEAIEYYQKALELDP NNAEAKQNLGNAKQKQGLEHHHHHH |
| HR04C4_1* | MHHHHHHGSWGSDECEEKARRVAEKVERLKRSGTSEDEIAEEVAREISEVI RTLKESGSSYEVICECVARIVAEIVEALKRSGTSAVEIAKIVARVISEVIRTLKE SGSSYEVICECVARIVAEIVEALKRSGTSAAIIALIVALVISEVIRTLKESGSSFE VILECVIRIVLEIIEALKRSGTSEQDVMLIVMAVLLVVLATLQLSGS |
| 2JFB (PDB ID)* | MAVKGLGEVDQKYDGSKLRIGILHARWNRKIIDALVAGAVKRLQEFGVKEEN IIIETVPGSFELPYGSKLFVEKQKRLGKPLDAIIPIGVLIKGSTMHFEYICDSTTH QLMKLNFELGIPVIFGVLTCLTDEQAEARAGLIEGKMHNHGEDWGAAAVEM ATKFN |
| 2OBX (PDB ID)* | MNQHSHKDYETVRIAVVRARWHADIVDQCVSAFEAEMADIGGDRFAVDVFD VPGAYEIPLHARTLAETGRYGAVLGTAFVVNGGIYRHEFVASAVIDGMMNVQ LSTGVPVLSAVLTPHNYHDSAEHHRFFFEHFTVKGKEAARACVEILAAREKI AA |
| 2B98 (PDB ID)* | MTKKVGIVDTTFARVDMASIAIKKLKELSPNIKIIRKTVPGIKDLPVACKKLLEE EGCDIVMALGMPGKAEKDKVCAHEASLGLMLAQLMTNKHIIEVFVHEDEAK DDKELDWLAKRRAEEHAENVYYLLFKPEYLTRMAGKGLRQGFEDAGPARE |

**Supplementary Table 7.** List of all designed homotrimers and pre-validated components tested with their corresponding amino acid sequences including initiating methionine and $His_6$-tag. Designs that expressed solubly are denoted in bold and under their name are the experimental methods used for characterization.
*Components from previously described designed homo-oligomers[1] or the Protein Data Bank (PDB IDs).

| Design | Sequence |
|---|---|
| T33_dn1A (1na0C3_3) | MGNLAEKMYKAGNAMYRKGQYTIAIIAYTLALLKDPNNAEAWYNLGNAAYKKGEYDE AIEAYQKALELDPNNAEAWYNLGNAYYKQGDYDEAIEYYERALELDPENAEAALNLLE AKEKQG |
| T33_dn2B (1na0C3_2) | MEEAELAYLLGELAYKLGEYRIAIRAYRIALKRDPNNAEAWYNLGNAYYKQGDYREAIK YYAKALTLDPKNAEAWYNLGNAVYKQGDYRIAILFYRAALKLDPNNAEAKQNLGNAKQ KQGLEHHHHHH |
| **T33_dn2A (1na0C3_3)** | **MGNLAEKMYKAGNAMYRKGQYTIAIIAYTLALLKDPNNAEAWYNLGNAAYKKGEYD EAIEAYQKALELDPNNAEAWYNLGNAYYKQGDYDEAIEYYKKALRLDPRNVDAIEN LIEAEEKQG** |
| **T33_dn2B (1na0C3_2)** | **MEEAELAYLLGELAYKLGEYRIAIRAYRIALKRDPNNAEAWYNLGNAYYKQGDYRE AIRYYLRALKLDPENAEAWYNLGNALYKQGKYDLAIIAYQAALEEDPNNAEAKQNL GNAKQKQGLEHHHHHH** |
| T33_dn3A (1na0C3_7) | MGNSAEAMYKMGNAAYKQGDYILAIIAYLLALEKDPNNAEAWYNLGNAAYKQGDYKE AILYYIRALQLDPNNAEAWYNLGNAFYKKGDYRVAIILYRMALKLDPNNAEAKQNLGNA KQKQGDIHHHHHH |
| T33_dn3B (1na0C3_2) | MEEAELAYLLGELAYKLGEYRIAIRAYRIALKRDPNNAEAWYNLGNAYYKQGDYDEAIE YYQKALELDPNNAEAWYNLGNAYYKQGDYEEAILYYLEALDLDPNNAEAAENLLNAV KKDE |
| **T33_dn4A (1na0C3_7)** | **MGNSAEAMYKMGNAAYKQGDYILAIIAYLLALEKDPNNAEAWYNLGNAAYKQGDY DEAIEYYQKALELDPNNAEAWYNLGNAYYKQGDYDEAIEYYEKALELDPRNAEALK NLLEAKAKQD** |
| **T33_dn4B (3ltjC3_1)** | **MHHHHTDPLAVILYIAILKAEKSIARAKAAEALGKIGDERAVEPLIKALKDEDALVRAA AADALGQIGDERAVEPLIKALKDEEGLVRASAAIALGQIGDERAVRPLIKALADERDL VRVAAAVALGRIGDERAVKPLIIVLLDEEGEVREAAAIALGSIGGERVRAAMEKLAER GRGFARKVAVNYLETHKLEHHHHHH** |
| **T33_dn5A 1na0C3_7** | **MGNSAEAMYKMGNAAYKQGDYILAIIAYLLALEKDPNNAEAWYNLGNAAYKQGDY DEAIEYYQKALELDPNNAEAWYNLGNAYYKQGDYDEAIEYYEKALELDPNNAEALK NLLEAIAEQD** |
| **T33_dn5B (3ltjC3_1)** | **MHHHHTDPLAVILYIAILKAEKSIARAKAAEALGKIGDERAVEPLIKALKDEDALVRAA AADALGQIGDERAVEPLIKALKDEEGLVRASAAIALGQIGDERAVQPLIKALTDERDL VRVAAAVALGRIGDEKAVRPLIIVLKDEEGEVREAAAIALGSIGGERVRAAMEKLAER GTGFARKVAVNYLETHKLEHHHHHH** |
| **T33_dn6A (1na0C3_2)** | **MGEEAELAYLLGELAYKLGEYRIAIRAYRIALDEDPDNAEAWYNLGNAYYKQGDYR EAILYYQMALRLDPNNAEAWYNLGNAYYKQGDYDRAIEYYQKALELDPNNAEAKQ NLGNAKQKQGDIHHHHHH** |
| **T33_dn6B (3ltjC3_1)** | **MHHHHTDPLAVILYIAILKAEKSIARAKAAEALGKIGDERAVEPLIKALKDEDALVRAA AADALGQIGDERAVEPLIKALKDEEGLVRASAAIALGQIGDKRAVRPLIRALKDERDL VREAAAVALGRIGDELAVEPLIKALKDEEGEVREAAAIALGSIGGEIVRMMMDKLAE TGTGFARKVAVNYLETHK** |

| | |
|---|---|
| **T33_dn7A<br>(1na0C3_2)** | **MGEEAELAYLLGELAYKLGEYRIAIRAYRIALKRDPNNAEAWYNLGNAYYKQGDYD<br>EAIEYYQKALELDPNNAEAWYNLGNAYYKQGDYDEAIEYYRKALELDPENEEALEN<br>LLNAKQKQGDIHHHHHH** |
| **T33_dn7B<br>(3ltjC3_1)** | **MHHHHTDPLAVILYIAILKAEKSIARAKAAEALGKIGDERAVEPLIKALKDEDALVRAA<br>AADALGQIGDERAVEPLIKALKDEEGLVRASAAIALGQIGDERAVEPLIKALKDERDL<br>VRVAAAVALGRIGDERAVEPLIKALKDEEGEVREAAAIALGSIGGKRVRLAMLKLAL<br>EGTGFARKVAVNYLETHK** |
| T33_dn8A<br>(1na0C3_2) | MGEEAELAYLLGELAYKLGEYRIAIRAYRIALKRDPNNAEAWYNLGNAYYKQGDYDEA<br>IEYYQKALELDPNNAEAWYNLGNAYYKQGDYDEAIEYYRKALELDPENLEALLNLLNA<br>KDKRG |
| T33_dn8B<br>(HR00C3_2) | MIEEVVAEMIDILAESSKKSIEELARAADNKTTEKAVAEAIEEIARLATAAIQLIEALAKNL<br>ASEEFMARAISAIAELAKKAIEAIYRLADNHTTDTFMARAIAAIANLAVTAILAIAALASNH<br>TTEEFMARAIRAIAELAKKAIEAIYRLADNHTTDKFMAAAIEAIALLATLAILAIALLASNHT<br>TERFMAKAILAIAVLAKKAIEAIYRLADNHTSPTYIEKAIEAIEKIARKAIKAIEMLAKNITTE<br>EYKEEAKSAIEIIRELARIAIRRLEDNRTLEHHHHHH |
| T33_dn9A<br>(1na0C3_2) | MGEEAELAYLLGELAYKLGEYRIAIRAYRIALKRDPNNAEAWYNLGNAYYKQGDYDEA<br>IEYYQKALELDPNNAEAWYNLGNAYYKQGDYDEAIEYYQKALELDPENLEAILNLGEA<br>KLKQG |
| T33_dn9B<br>(HR00C3_2) | MIEEVVAEMIDILAESSKKSIEELARAADNKTTEKAVAEAIEEIARLATAAIQLIEALAKNL<br>ASEEFMARAISAIAELAKKAIEAIYRLADNHTTDTFMARAIAAIANLAVTAILAIAALASNH<br>TTEEFMARAISAIAELAKKAIEAIYRLADNHTTDKFMAAAIEAIALLATLAILAIALLASNHT<br>TEKFMAEAIIVIALLAVLAIMAIYRLADNHTSPTYIEKAIEAIEKIARKAIKAIEMLAKNITTEE<br>YKEKAKSAIDLIRQLADIIIRKLEDNRTLEHHHHHH |
| **T33_dn10A<br>(1na0C3_2)** | **MGEEAELAYLLGELAYKLGEYRIAIRAYRIALKRDPNNAEAWYNLGNAYYKQGDYD<br>EAIEYYQKALELDPNNAEAWYNLGNAYYKQGDYDEAIEYYEKALELDPENLEALQN<br>LLNAMDKQG** |
| **T33_dn10B<br>(HR00C3_2)** | **MIEEVVAEMIDILAESSKKSIEELARAADNKTTEKAVAEAIEEIARLATAAIQLIEALAK<br>NLASEEFMARAISAIAELAKKAIEAIYRLADNHTTDTFMARAIAAIANLAVTAILAIAAL<br>ASNHTTEEFMARAISAIAELAKKAIEAIYRLADNHTTDKFMAAAIEAIALLATLAILAIA<br>LLASNHTTEKFMARAIMAIAILAAKAIEAIYRLADNHTSPTYIEKAIEAIEKIARKAIKAIE<br>MLAKNITTEEYKEKAKKIIDIIRKLAKMAIKKLEDNRTLEHHHHHH** |
| T33_dn11A<br>(1na0C3_2) | MGEEAELAYLLGELAYKLGEYRIAIRAYRIALKRDPNNAEAWYNLGNAYYKQGDYDEA<br>IEYYQKALELDPNNAEAWYNLGNAYYKQGDYDEAIEYYRKALELDKENIEALLNLLNAK<br>EKQD |
| T33_dn11B<br>(HR00C3_2) | MIEEVVAEMIDILAESSKKSIEELARAADNKTTEKAVAEAIEEIARLATAAIQLIEALAKNL<br>ASEEFMARAISAIAELAKKAIEAIYRLADNHTTDTFMARAIAAIANLAVTAILAIAALASNH<br>TTEEFMARAISAIAELAKKAIEAIYRLADNHTTDKFMAAAIEAIALLATLAILAIALLASNHT<br>TERFMAKAILAIAILAAKAIEAIYRLADNHTSPTYIEKAIEAIEKIARKAIKAIEMLAKNITTEE<br>YKEEAKSAIEIIRLLAKAVIKRLQDNRTLEHHHHHH |
| **O32_dn1A<br>(1na0C3_3)** | **MGELAEKMYKAGNAMYRKGQYTIAIIAYTLALLKDPNNAEAWYNLGNAAYKKGEYD<br>EAIVAYVEALELDPNNAEAWYNLGNAYYKQGDYEEAIEYYQKALELDPNNAEAKQN<br>LGNAKQKQG** |
| **O32_dn1B<br>(ank1C2_1)** | |

| | |
|---|---|
| | **MSRRGRLLIIAAENGNKDRVKDLIQRGADVNASDRRGRTPLHHAAENGHAEVVALLI<br>EKGADVNAKDSDGRTPLHHAAENGHDEVVLILLLKGADVNAKDSDGRTPLHHAAE<br>NGHKRVVLVLILAGADVNTSDSDGRTPLDLAREHGNEEVVKALEKQLEHHHHHH** |
| O32_dn2A<br>(1na0C3_2) | MGEEAELAYLLGELAYKLGEYRIAIRAYRIALKRDPNNAEAWYNLGNAYYKQGDYDEA<br>IEYYQKALELDPNNAEAWYNLGNAYYKQGDYDEAIEYYQKALELDPNNAEARKNLIIA<br>DLKQEDIHHHHHH |
| O32_dn2B<br>(ank1C2_1) | MSELGEALILAAERGKKDRVKDLIEEGADVNASDSDGRTPLHHAAENGHAEVVALLIE<br>KGADVNAKDSDGRTPLHHAAENGHDEVVLILLLKGADVNAKDSDGRTPLHHAAENGH<br>KRVVLVLILAGADVNTSDSDGRTPLDLAREHGNEEVVKALEKQ |
| **O32_dn3A<br>(3ltjC3_1)** | **MGHHHHHHGWHHHHTDPLAVILYIAILKAEKSIARAKAAEALGKIGDERAVEPLIKAL<br>KDEDALVRAAAADALGQIGDERAVEPLIEALEDEEGLVRASAAIALGQIGDERAVEP<br>LILALADERDLVRVAAAVALGRIGDERAVEPLIVMLRDEEGEVREAAAIALGSIGGER<br>VRAAMEELAERGRGFARKVAVNYLETHK** |
| **O32_dn3B<br>(ank1C2_1)** | **MSELGKRLIEAAENGNKKRVKDLIENGADVNASDSDGRTPLHHAAENGHAEVVALL<br>IEKGADVNAKDSDGRTPLHHAAENGHDEVVLILLLKGADVNAKDSDGRTPLHHAAE<br>NGHKRVVLVLILAGADVNTKDEEGDTPLALALEHGNREVIKALLKQ** |
| O43_dn1A<br>(1na0C4_1) | MGTLARVAYILGAIAYAQGEYDIAITAYQVALDLDPNNAEAWYNLGNAYYKQGDYDEAI<br>EYYQKALELDPNNAEAWYNLGNAYYKQGDYLLAIVYYAKALILDPNNAEAKQNLGNAI<br>QKQD |
| O43_dn1B<br>(1na0C3_3) | MNLAEKMYKAGNAMYRKGQYTIAIIAYTLALLKDPNNAEAWYNLGNAAYKKGEYDEAI<br>EAYQKALELDPNNAEAWYNLGNAYYKQGDYLEAIAYYAKALLLDPNNAEARQNLGNA<br>MQKSELEHHHHHH |
| O43_dn2A<br>(1na0C4_1) | MGTLARVAYILGAIAYAQGEYDIAITAYQVALDLDPNNAEAWYNLGNAYYKQGDYDEAI<br>EYYQKALELDPNNAEAWYNLGNAYYKQGDYDEAILYYVKALVLDPNNAEAKQNLGNA<br>RQKQG |
| O43_dn2B<br>(1na0C3_3) | MNLAEKMYKAGNAMYRKGQYTIAIIAYTLALLKDPNNAEAWYNLGNAAYKKGEYDEAI<br>EAYQKALELDPNNAEAWYNLGNAYYKQGDYLEAILYYVKALKLDPNNAEAKQNLGNA<br>EQKKDLEHHHHHH |
| O43_dn3A<br>(1na0C4_1) | MGTLARVAYILGAIAYAQGEYDIAITAYQVALDLDPNNAEAWYNLGNAYYKQGDYDEAI<br>KYYQKALELDPNNAEAWYNLGNAYYKQGDYVIAIALYQLALELDPNNAEAKQNLGNA<br>EQKEGDIHHHHHH |
| O43_dn3B<br>(1na0C3_7) | MNSAEAMYKMGNAAYKQGDYILAIIAYLLALEKDPNNAEAWYNLGNAAYKQGDYDEAI<br>EYYQKALELDPNNAEAWYNLGNAYYKQGDYLEAIEYYIKALELDPNNEEARQNLLNAA<br>KKIE |
| **O43_dn4A<br>(1na0C4_1)** | **MGTLARVAYILGAIAYAQGEYDIAITAYQVALDLDPNNAEAWYNLGNAYYKQGDYDE<br>AIEYYQKALELDPNNAEAWYNLGNAYYKQGDYEEAIEYYLKALELDPNNAEARQNL<br>RNAMQKEG** |
| **O43_dn4B<br>(1na0C3_7)** | **MNSAEAMYKMGNAAYKQGDYILAIIAYLLALEKDPNNAEAWYNLGNAAYKQGDYD<br>EAIEYYQKALELDPNNAEAWYNLGNAYYKQGDYLAAIIYYRRALELDPNNAEAKQN<br>LGNAEQKEGLEHHHHHH** |

| | |
|---|---|
| O43_dn5A<br>(1na0C4_1) | MGTLARVAYILGAIAYAQGEYDIAITAYQVALDLDPNNAEAWYNLGNAYYKQGDYDEAIEYYQKALELDPNNAEAWYNLGNAYYKQGDYREALRYYIKALKLDPNNAEAKQNLGNALEKRG |
| O43_dn5B<br>(1na0C3_7) | MNSAEAMYKMGNAAYKQGDYILAIIAYLLALEKDPNNAEAWYNLGNAAYKQGDYDEAIEYYQKALELDPNNAEAWYNLGNAYYKQGDYLVAIIYYLEALELDPNNAEAKQNLGNAKQKEGLEHHHHHH |
| **O43_dn6A<br>(1na0C4_1)** | **MGTLARVAYILGAIAYAQGEYDIAITAYQVALDLDPNNAEAWYNLGNAYYKQGDYDEAIEYYQKALELDPNNAEAWYNLGNAYYKQGDYQEAIEYYARALRRDRRNKEAIENLINALQKED** |
| **O43_dn6B<br>(1na0C3_2)** | **MEEAELAYLLGELAYKLGEYRIAIRAYRIALKRDPNNAEAWYNLGNAYYKQGRYVRALIYYLRALLLDPENAEAWYNLGNAYYKKGDYDIAIVYYELALEDDPNNAEAKQLLGNAKQKQGLEHHHHHH** |
| O43_dn7A<br>(1na0C4_1) | MGTLARVAYILGAIAYAQGEYDIAITAYQVALDLDPNNAEAWYNLGNAYYKQGDYDEAIEYYQKALELDPNNAEAWYNLGNAYYKQGDYDEAIEYYKKALRLDPNNEEAKQNLMNALQKQD |
| O43_dn7B<br>(3ltjC3_1) | MHHHHTDPLAVILYIAILKAEKSIARAKAAEALGKIGDERAVEPLIKALKDEDALVRAAAADALGQIGDERAVEPLIKALKDEEGLVRASAAIALGQIGDERAVEPLIKALKDERDLVRVAAAVALGRIGDKKAVLPLIKALKDEEGEVREAAAIALGSIGGRLVRAMMELLAETGRGFARKVAVNYLETHKLEHHHHHH |
| O43_dn8A<br>(HR04C4_1) | MGDECEEKARLLAELVETLKRSGTSEDEIAEDVARLISEMIRNLKESGSSYEVICECVARIVAEIVEALKRSGTSAVEIAKIVARVISEVIRTLKESGSSYEVICECVARIVAEIVEALKRSGTSAAIIALIVALVISEVIRTLKESGSSFEVILECVIRIVLEIIEALKRSGTSEQDVMLIVMAVLLVVLATLQLSGS |
| O43_dn8B<br>(1na0C3_3) | MNLAEKMYKAGNAMYRKGQYTIAIIAYTLALLKDPNNAEAWYNLGNAAYKKGEYDEAIEAYQKALELEPNNAEAWYNLGNAYYKQGDYEEAIIYYLKALVLDPRNAEARQNLGNAKQKEGLEHHHHHH |
| O43_dn9A<br>(HR04C4_1) | MGDECEELARIVAELVEKLKRSGTSEDEIAERVAREISEVIKLLKKSGSSYEVICECVARIVAEIVEALKRSGTSAVEIAKIVARVISEVIRTLKESGSSYEVICECVARIVAEIVEALKRSGTSAAIIALIVALVISEVIRTLKESGSSFEVILECVIRIVLEIIEALKRSGTSEQDVMLIVMAVLLVVLATLQLSGS |
| O43_dn9B<br>(1na0C3_3) | MNLAEKMYKAGNAMYRKGQYTIAIIAYTLALLKDPNNAEAWYNLGNAAYKKGEYDEAIEAYQKALELDPENAEAWYNLGNAYYKQGEYLEALLYYLKALILDPNNAEAKQNLGNARQKQGLEHHHHHH |
| O43_dn10A<br>(HR04C4_1) | MGDECERKARLVAKIVELLKRSGTSEDEIAEEVARLISLVIKVLKKSGSSYEVICECVARIVAEIVEALKRSGTSAVEIAKIVARVISEVIRTLKESGSSYEVICECVARIVAEIVEALKRSGTSAAIIALIVALVISEVIRTLKESGSSFEVILECVIRIVLEIIEALKRSGTSEQDVMLIVMAVLLVVLATLQLSGS |
| O43_dn10B<br>(1na0C3_3) | MNLAEKMYKAGNAMYRKGQYTIAIIAYTLALLKDPNNAEAWYNLGNAAYKKGEYDEAIEAYQKALELDPENAEAWYNLGNAYYKQGDYAEAMLYYLKALLLDPNNAEAKQNLGNAEQKAGLEHHHHHH |
| O43_dn11A<br>(HR04C4_1) | MGDECEEKAELVALLVEALKKLGTSEDEIAEEVAKEISRVIRRLKESGSSYEVICECVARIVAEIVEALKRSGTSAVEIAKIVARVISEVIRTLKESGSSYEVICECVARIVAEIVEALKR |

| | |
|---|---|
| | SGTSAAIIALIVALVISEVIRTLKESGSSFEVILECVIRIVLEIIEALKRSGTSEQDVMLIVMA<br>VLLVVLATLQLSGS |
| O43_dn11B (1na0C3_7) | MNSAEAMYKMGNAAYKQGDYILAIIAYLLALEKDPNNAEAWYNLGNAAYKQGDYDEAI<br>EYYQKALELDPENAEAWYNLGNAYYKQGDYELAIIFYKVALALDPNNAEAKQNLGNAK<br>QKQGLEHHHHHH |
| **O43_dn12A (HR04C4_1)** | **MGDRCERRAKLVALKVELLKKDGTSEDEIAEEVAREISEVIRDLRKSGSSYEVICECV<br>ARIVAEIVEALKRSGTSAVEIAKIVARVISEVIRTLKESGSSYEVICECVARIVAEIVEAL<br>KRSGTSAAIIALIVALVISEVIRTLKESGSSFEVILECVIRIVLEIIEALKRSGTSEQDVMLI<br>VMAVLLVVLATLQLSGS** |
| **O43_dn12B (1na0C3_2)** | **MEEAELAYLLGELAYKLGEYRIAIRAYRIALKRDPNNAEAWYNLGNAYYKQGDYDE<br>AIEYYQKALELDPNNAEAWYNLGNAYYKQGDYDEAIEYYQKALELDPNNIKAELNLII<br>AEEKQGLEHHHHHH** |
| O43_dn13A (HR04C4_1) | MGDECEELARAVALVVEILKRSGTSEDEIAEEVARLISRVIRKLKESGSSYEVICECVAR<br>IVAEIVEALKRSGTSAVEIAKIVARVISEVIRTLKESGSSYEVICECVARIVAEIVEALKRS<br>GTSAAIIALIVALVISEVIRTLKESGSSFEVILECVIRIVLEIIEALKRSGTSEQDVMLIVMAV<br>LLVVLATLQLSGS |
| O43_dn13B (1na0C3_2) | MEEAELAYLLGELAYKLGEYRIAIRAYRIALKRDPNNAEAWYNLGNAYYKQGDYDEAIE<br>YYQKALELDPNNAEAWYNLGNAYYKKGDYLIAILYYLVALTLDPNNAEAKQNLGNAKQ<br>KDGLEHHHHHH |
| O43_dn14A (HR04C4_1) | MGDKCEEMAELVAQLVELLKESGTSEDEIAEKVARLISKVIRKLKESGSSYEVICECVA<br>RIVAEIVEALKRSGTSAVEIAKIVARVISEVIRTLKESGSSYEVICECVARIVAEIVEALKR<br>SGTSAAIIALIVALVISEVIRTLKESGSSFEVILECVIRIVLEIIEALKRSGTSEQDVMLIVMA<br>VLLVVLATLQLSGS |
| O43_dn14B (1na0C3_2) | MEEAELAYLLGELAYKLGEYRIAIRAYRIALKRDPNNAEAWYNLGNAYYKQGDYDEAIE<br>YYMKALKLDPKNAEAWYNLGNAYYKQGDYLLAILIYEMALILDPNNAEAKQNLGNAKQ<br>KEGLEHHHHHH |
| O43_dn15A (HR04C4_1) | MGRKCELLARLVAMIVELLKESGTSEDEIAEEVAREISEVIRTLKEEGSSYEVICECVAR<br>IVAEIVEALKRSGTSAVEIAKIVARVISEVIRTLKESGSSYEVICECVARIVAEIVEALKRS<br>GTSAAIIALIVALVISEVIRTLKESGSSFEVILECVIRIVLEIIEALKRSGTSEQDVMLIVMAV<br>LLVVLATLQLSGS |
| O43_dn15B (1na0C3_2) | MEEAELAYLLGELAYKLGEYRIAIRAYRIALKRDPNNAEAWYNLGNAYYKQGDYDEAIE<br>YYQKALELDPRNAEAWYNLGNAYYKQGDYLMAILIYQLALMLDPNNAEAKQNLGNAK<br>QKRGLEHHHHHH |
| O43_dn16A (HR04C4_1) | MGEDCEELAELVAELVERLKRRGTSEDEIAEEVARIISEVIRMLKESGSSYEVICECVA<br>RIVAEIVEALKRSGTSAVEIAKIVARVISEVIRTLKESGSSYEVICECVARIVAEIVEALKR<br>SGTSAAIIALIVALVISEVIRTLKESGSSFEVILECVIRIVLEIIEALKRSGTSEQDVMLIVMA<br>VLLVVLATLQLSGS |
| O43_dn16B (1na0C3_2) | MEEAELAYLLGELAYKLGEYRIAIRAYRIALKRDPNNAEAWYNLGNAYYKQGDYKEAIK<br>YYQKALKLDPNNAEAWYNLGNAYYKKGDYIMAILAYELALEEDPNNAEAKQNLGNAK<br>QKQGLEHHHHHH |
| **O43_dn17A (HR04C4_1)** | **MGDECEEKARRVALKVLKLRLRGTSEDEIAEEVAREISKVIETLKESGSSYEVICECV<br>ARIVAEIVEALKRSGTSAVEIAKIVARVISEVIRTLKESGSSYEVICECVARIVAEIVEAL** |

| | |
|---|---|
| | **KRSGTSAAIIALIVALVISEVIRTLKESGSSFEVILECVIRIVLEIIEALKRSGTSEQDVMLI<br>VMAVLLVVLATLQLSGS** |
| **O43_dn17B<br>(1na0C3_2)** | **MEEAELAYLLGELAYKLGEYRIAIRAYRIALKRDPNNAEAWYNLGNAYYKQGDYDE<br>AIEYYQKALELDPNNAEAWYNLGNAYYKQGDYDEAIEYYQKALELDPNNEEAKIVL<br>GLAKEEQELEHHHHHH** |
| **O43_dn18A<br>(HR04C4_1)** | **MDRCEELARRIAEVVERAKRAGTSEDEIAESVARVISLVIRALKLSGSSYEVICECVA<br>RIVAEIVEALKRSGTSAVEIAKIVARVISEVIRTLKESGSSYEVICECVARIVAEIVEALK<br>RSGTSAAIIALIVALVISEVIRTLKESGSSFEVILECVIRIVLEIIEALKRSGTSEQDVMLIV<br>MAVLLVVLATLQLSGSGGWLEHHHHHH** |
| **O43_dn18B<br>(1na0C3_2)** | **MGEEAELAYLLGELAYKLGEYRIAIRAYRIALKRDPNNAEAWYNLGNAYYKQGDYD<br>EAIEYYQKALELDPNNAEAWYNLGNAYYKQGDYDEAIEYYQKALELDPSNLDAAVN<br>LGAATMLTS** |
| **I32_dn1A<br>(1na0C3_3)** | **MGNLAEKMYKAGNAMYRKGQYTIAIIAYTLALLKDPNNAEAWYNLGNAAYKKGEYD<br>EAIEAYQKALELEPENAEALYNLGNAYYKQGEYDEAILYYLIALELDPNNAEAKQNL<br>GNAKQKQGDIHHHHHH** |
| **I32_dn1B<br>(ank1C2_1)** | **MSRLGIRLIIAAIEGNKDRVKDLIENGADVNASDSVGRTPLHHAAENGHAEVVALLIE<br>KGADVNAKDSDGRTPLHHAAENGHDEVVLILLLKGADVNAKDRDGRTPLHHAAEN<br>GHKRVVLVLILAGADVNTSDSDGRTPLDLAREHGNEEVVKALEKQ** |
| **I32_dn2A<br>(1na0C3_3)** | **MGDLAEKMYKAGNAMYRKGQYTIAIIAYTLALLKDPNNAEAWYNLGNAAYKKGEYD<br>EAILAYLKALELDPNNAEAWYNLGNAFYKQGDYRMAIKYYQKALELDPNNAEAKQN<br>LGNAKQKQG** |
| **I32_dn2B<br>(ank1C2_1)** | **MSELGELLIVAAENGNKKMVRDLIKNGADVNASDEDGRTPLHHAAENGHAEVVALL<br>IEKGADVNAKDSDGRTPLHHAAENGHDEVVLILLLKGADVNAKDSDGRTPLHHAAE<br>NGHKRVVLVLILAGADVNTSDSDGRTPLDLAREHGNEEVVKALEKQLEHHHHHH** |
| I32_dn3A<br>(1na0C3_3) | MGRLAKKMYKAGNAMYRKGQYTIAIIAYTLALLKDPNNAEAWYNLGNAAYKKGEYAE<br>AIVAYIKALELDPNNAEAWYNLGNALYKLGAYNAAIQVYQKALELDPNNAEAKQNLGN<br>AKQKKG |
| I32_dn3B<br>(ank1C2_1) | MKILGLALIAAARNGEKERVETLIEAGADVNASDDDGRTPLHHAAENGHAEVVALLIEK<br>GADVNAKDSDGRTPLHHAAENGHDEVVLILLLKGADVNAKDSDGRTPLHHAAENGHK<br>RVVLVLILAGADVNTSDSDGRTPLDLAREHGNEEVVKALEKQLEHHHHHH |
| I32_dn4A<br>(1na0C3_7) | MGKSAEAMYKMGNAAYKQGDYILAIIAYLLALEKDPKNAEAWYNLGNAAYKQGDYEE<br>AIRYYLKALLLDDNNAEAWYNLGNAYYKQGDYREAIMLYQKALELDPNNAEAKQNLG<br>NAKQKQG |
| I32_dn4B<br>(ank1C2_1) | MSELGKLLIMAAELGNKRLVKELIENGADVNASDSDGRTPLHHAAENGHAEVVALLIE<br>KGADVNAKDSDGRTPLHHAAEEGHDEVVLILLLKGADVNAKDSDGRTPLHHAAENGH<br>KRVVLVLILAGADVNTSDSDGRTPLDLAREHGNEEVVKALEKQLEHHHHHH |
| I32_dn5A<br>(1na0C3_7) | MGRSAEAMYKMGNAAYKQGDYILAIIAYLLALEKDPNNAEAWYNLGNAAYKQGDYRE<br>AIRYYLKALALDPNNAEAWYNLGNAFYKQGDYNEAIEVYQKALELDPNNAEAKQNLG<br>NAKQKQG |
| I32_dn5B<br>(ank1C2_1) | MSELGRMLIEAAELGKKEIVKELIENGADVNASDSDGRTPLHHAAENGHAEVVALLIEK<br>GADVNAKDSDGRTPLHHAAENGHDEVVLILLLKGADVNAKDSDGRTPLHHAAENGHK<br>RVVLVLILAGADVNTSDSDGRTPLDLAREHGNEEVVKALEKQLEHHHHHH |

| | |
|---|---|
| **I32_dn6A (1na0C3_2)** | **MGEEAELAYLLGELAYKLGEYRIAIRAYRIALKRDPNNAEAWYNLGNAYYKQGDYDEAIEYYQKALELDPNNAEAWYNLGNAYYKQGDYDEAIEYYQKALELDPNNDEADDNLLNADQKQDDIHHHHHH** |
| **I32_dn6B (ank1C2_1)** | **MSRLGKKLIIAAERGNKDRVKDLIENGADVNASDEDGRTPLHHAAENGHAEVVALLIEKGADVNAKDSDGRTPLHHAAENGHDEVVLILLLKGADVNAKDSDGRTPLHHAAENGHKRVVLVLILAGADVNTSDSDGRTPLDLAREHGNEEVVKALEKQ** |
| **I32_dn7A (1na0C3_2)** | **MGEEAELAYLLGELAYKLGEYRIAIRAYRIALKRDPNNAEAWYNLGNAYYKQGDYDEAIEYYQKALELDPNNAEAWYNLGNAYYKQGDYDEAIEYYQKALELDPKNMEALLDLGNAKQKQKDIHHHHHH** |
| **I32_dn7B (ank1C2_1)** | **MSELGKDLIVAAALGNKDRVKDLIENGADVNASDRRGATPLHMAALNGHAEVVALLIEKGADVNAKDSDGRTPLHHAAENGHDEVVLILLLKGADVNAKDSDGRTPLHHAAENGHKRVVLVLILAGADVNTSDSDGRTPLDLAREHGNEEVVKALEKQ** |
| **I32_dn8A (1na0C3_2)** | **MGEEAELAYLLGELAYKLGEYRIAIRAYRIALKRDPNNAEAWYNLGNAYYKQGDYDEAIEYYQKALELDPNNAEAWYNLGNAYYKQGDYLRAIAYYRKALELDPNNAEAKQNLGNAKQKIE** |
| **I32_dn8B (ank3C2_1)** | **MELEGERLIEAAENGNKDRVKDLLENGALVNASDSDGKTPLHLAAENGHAKVVLLLLEQGAKPNAKDSDGKTPLHLAAENGHAVVVALLLMHGADPNAKDSDGKTPLHLAAENGHEEVVILLLAMGADPNTSDSDGRTPLDLAREHGNEEVVKVLEDHGGWLEHHHHALEHHHHHH** |
| I32_dn9A (HR00C3_2) | MGIEEVVAEMIDILAESSKKSIEELARAADNKTTEKAVAEAIEEIARLATAAIQLIEALAKNLASEEFMADAISAIAELAKKAIEAIYRLADNHTTDTFMARAIAAIANLAVTAILAIAALASNHTTEQFMAIAIIAIAELAKKAIEAIYRLADNHTTDKFMAAAIEAIALLATLAILAIALLASNHTTEAFMALAILLIAELAKKAIEAIYRLADNHTSPTYIEKAIEAIEKIARKAIKAIEMLAKNITTEEYKEKARAAILEIREKAKEAIKRLEDNRT |
| I32_dn9B (ank1C2_1) | MHHHHHHSELGKRLIEAAENGNKKRVLELIENGADVNASDSDGRTPLHHAAENGHAEVVALLIELGADVNAKDSDGRTPLHHAAENGHDEVVLILLLKGADVNAKDSDGRTPLHHAAENGHKRVVLVLILAGADVETSDSRGRTPLMLAVEHGNIEVALALLKQGW |
| **I32_dn10A (HR00C3_2)** | **MGHHHHHHWGIEEVVAEMIDILAESSKKSIEELARAADNKTTEKAVAEAIEEIARLATAAIQLIEALAKNLASEEFMARAISAIAELAKKAIEAIYRLADNHTTDTFMARAIAAIANLAVTAILAIAALASNHTTEEFMARAISAIAELAKKAIEAIYRLADNHTTDKFMAAAIEAIALLATLAILAIALLASNHTTEEFMAKAISAIARLAKKAILAIYKLADNHTSPTYIEKAIEAIEKIARKAIKAIEMLAKNITTEEYKEKAKSAIDEIREIAKIAIKTLEDNRT** |
| **I32_dn10B (ank1C2_1)** | **MSEIGKRLIEAAENGNKERVKLLIELGADVNASDSDGRTPLHHAAENGHAEVVALLIEKGADVNAKDSDGRTPLHHAAENGHDEVVLILLLKGADVNAKDSDGRTPLHHAAENGHKRVVLVLILAGADVNTSDSDGRTPLDLAREHGNEEVVKALEKQ** |
| I53_dn1A (2B98) | MGHHHHHHKKVGIVDTTFARVDMAIMAIIVLELRPRNIKIIRKTVPGIKDLPVACKKLLEEEGCDIVMALGMPGKAEKDKVCAHEASLGLMLAQLMTNKHIIEVFVHEDEAKDDRELDWLAKRRAEEHAENVYYLLFKPEYLTEMAGKGLRQGFEDAGP |
| I53_dn1B (HR00C3_2) | MIEEVVAEMIDILAESSKKSIRELAKAAKNKTTEKAVAEAIEEIARLATAAIQLIEALAKNLASEEFMARAISAIAELAKKAIEAIYRLADKHKTDTFMARAIAAIANLAVTAILAIAALASNHTTEEFMARAISAIAELAKKAIMAILLLALLHTTDKFMAAAIEAIALLATLAILAIALLASNHTTEEFMAKAISAIAELAKKAIEAIYLLADLHTPVLYEDKAIEAIEKIARKAIKAIEMLAKNITTEEYKEKAKSAIDEIREKAKEAIKRLERNRE |

| | |
|---|---|
| I53_dn2A (2JFB) | MGRYDGSKLRIGILHARWNRSIILALVLGAIERLLEFGVRAKNIIIETVPGSFELPYGSKLFVEKQKRLGKPLDAIIPIGVLIKGSTMHFEYICDSTTHQLMKLNFELGIPVIFGVLTCLTDEQAEARAGLIDGKMHNHGEDWGAAAVEMATKFN |
| I53_dn2B (1na0C3_2) | MEEAELAYLLGELAYKLGEYRIAIRAYRIALKRDPNNAEAWYNLGNAYYKQGDYDEAIEYYRRALKLEPENAEAWYNLGNAYYKQGDYKEAIAYYLIALILDPNNAEAKQNLGNAEQKQDLEHHHHHH |
| I53_dn3A (2JFB) | MGKYDGSKLRIGILHARWNRAIIIALVLGALKRLLEFGVKAKNIIIETVPGSFELPYGSKLFVEKQKRLGKPLDAIIPIGVLIKGSTMHFEYICDSTTHQLMKLNFELGIPVIFGVLTCLTDEQAEARAGLIKGKMHNHGEDWGAAAVEMATKFN |
| I53_dn3B (1na0C3_2) | MEEAELAYLLGELAYKLGEYRIAIRAYRIALKRDPNNAEAWYNLGNAYYKQGDYDEAIEYYQEALELDPENAEAWYNLGNAYYKQGDYKEALAYYLLALELDPNNAEAEQNLGNAEQKRDLEHHHHHH |
| I53_dn4A (2JFB) | MGKYDGSKLRIGILHARWNVKIIIALILGAIKRLREFGVKRENIIIEIVPGSFELPYGSKLFVEKQKRLGKPLDAIIPIGVLIKGSTMHFEYICDSTTHQLMKLNFELGIPVIFGVLTCLTDEQAEARAGLIEGKMHNHGEDWGAAAVEMATKFN |
| I53_dn4B (1na0C3_2) | MEEAELAYLLGELAYKLGEYRIAIRAYRIALKRDPNNAEAWYNLGNAYYKQGDYDEAIEYYQKALELDPNNAEAWYNLGNAYYKQGDYDEAIEYYQKALELDPNNLDAVMNLLEASLKQELEHHHHHH |
| **I53_dn5A (2JFB)** | **MGKYDGSKLRIGILHARWNAEIILALVLGALKRLQEFGVKRENIIIETVPGSFELPYGSKLFVEKQKRLGKPLDAIIPIGVLIKGSTMHFEYICDSTTHQLMKLNFELGIPVIFGVLTCLTDEQAEARAGLIEGKMHNHGEDWGAAAVEMATKFN** |
| **I53_dn5B (1na0C3_2)** | **MEEAELAYLLGELAYKLGEYRIAIRAYRIALKRDPNNAEAWYNLGNAYYKQGRYREAIEYYQKALELDPNNAEAWYNLGNAYYERGEYEEAIEYYRKALRLDPNNADAMQNLLNAKMREELEHHHHHH** |
| **I53_dn6A (2JFB)** | **MGDYDGSKLRIGILHARKNTEIIVALVIGAVERLEEFGVKRENIIIEIVPGSFELPYGSKLFVEKQKRLGKPLDAIIPIGVLIKGSTMHFEYICDSTTHQLMKLNFELGIPVIFGVLTCLTDEQAEARAGLIEGKMHNHGEDWGAAAVEMATKFN** |
| **I53_dn6B (1na0C3_2)** | **MEEAELAYLLGELAYKLGEYRIAIRAYRIALKRDPNNAEAWYNLGNAYYKQGDYDEAIEYYQKALELDPNNAEAWYNLGNAYYKQGDYDEAIEYYKKALRLDPDNAKALLNLIEAILKQKLEHHHHHH** |
| I53_dn7A (2JFB) | MGKYDGSKLRIGILHARWNRRIILALVIGAIIRLLEFGVKEDNIIIETVPGSFELPYGSKLFVEKQKRLGKPLDAIIPIGVLIKGSTMHFEYICDSTTHQLMKLNFELGIPVIFGVLTCLTDEQAEARAGLIEGKMHNHGEDWGAAAVEMATKFN |
| I53_dn7B (3ltjC3_1) | MHHHHTDPLAVILYIAILKAEKSIARAKAAEALGKIGDERAVEPLIKALKDEDALVRAAAADALGQIGDERAVIPLLRALLDKEGLVRASAAIALGQIGDKRAVLILILALEDERDLVRVAAAVALGRIGDEKAVEPLIEALKDEEGEVREAAAIALGSIGGERVRAAMEKLAETGTGFARKVAVNYLETHKLEHHHHHH |
| **I53_dn8A (2OBX)** | **MGHHHHHHHKDYETVRIAVVRARWHADIVRQCVMAFMKEMMRIGGRRFAVEVFDVPGAYEIPLHARTLAETGRYGAVLGTAFVVNGGIYRHEFVASAVIDGMMNVQLSTGVPVLSAVLTPHNYHDSAEHHRFFFEHFTVKGKEAARACVEILAARERI** |
| **I53_dn8B (1na0C3_2)** | |

| | |
|---|---|
| | **MEEAELAYLLGELAYKLGEYRIAIRAYRIALKRDPNNAEAWYNLGNAYYKQGDYDE AIEYYQKALELDPNNAEAWYNLGNAYYKQGDYDEAIEYYQKALELDPENEEAIDNLL EARQKQE** |
| **I53_dn9A (2OBX)** | **MGHHHHHHHKDYETVRIAVVRARWHAEIVDVCVLAFEIEMLDIGGDRFAVDVFDVPG AYEIPLHARTLAETGRYGAVLGTAFVVNGGIYRHEFVASAVIDGMMNVQLSTGVPVL SAVLTPHNYHDSAEHHEFFFEHFTVKGKEAARACVEILAAREKI** |
| **I53_dn9B (HR00C3_2)** | **MIEEVVAEMIDILAESSKKSIEELARAADNKTTEKAVAEAIEEIARLATAAIQLIEALAK NLASEEFMARAISAIAELAKKAIEAIYRLADNHTTDTFMARAIAAIANLAVTAILAIAAL ASNHTTEEFMARAISAIAELAKKAIAAIYRLADNHKTDKFMAAAIEAIALLATLAILAIA LLASNHTTEEFMAKAIRAIAKLAKMAILVIYALAIMHTSPTYIEKAIEAIEKIARKAIKAIE MLAKNITTEEYKEKAKSAIDEIREKAKEAIKRLEDKRE** |
| **I53_dn10A (2OBX)** | **MGHHHHHHHKDYETVRIAVVRARWHADIVDLCVIAFELEMLLIGGRRFAVDVFDVPG AYEIPLHARTLAETGRYGAVLGTAFVVNGGIYRHEFVASAVIDGMMNVQLSTGVPVL SAVLTPHNYHDSKRHHRFFAMHFIKKGKEAARACVEILAAREKI** |
| **I53_dn10B (HR00C3_2)** | **MIEEVVAEMIDILAESSKKSIEELAKAADNKTTEKAVAEAIEEIARLATAAIQLIEALAK NLASEEFMARAISAIAELAKKAIEAIYRLADNHTTDTFMARAIAAIANLAVTAILAIAAL ASNHTTEEFMARAISAIAELAKKAIEAILELALEHETDKFMAAAIEAIALLATLAILAIAL LASNHTTEEFMAKAIEAIAQLAKLAIIAIYLLALLHESPTYIEKAIEAIEKIARKAIKAIEM LAKNITTEEYKEKAKSAIDEIREKAKEAIKRLEDKRE** |
| I53_dn11A (2OBX) | MGHHHHHHHKDYETVRIAVVRARWHADIVDQCVSAFEREMAKIGGDRFAVDVFDVP GAYEIPLHARTLAETGRYGAVLGTAFVVNGGIYRHEFVASAVIDGMMNVQLSTGVPVL SAVLTPHEYHDSEIHHKIFFLLFTEKGKEAARACVEILAAREKI |
| I53_dn11B (HR00C3_2) | MIEEVVAEMIDILAESSKKSIEELARAADNKTTEKAVAEAIEEIARLATAAIQLIEALAKNL ASEEFMARAISAIAELAKKAIEAIYRLADNHTTDTFMARAIAAIANLAVTAILAIAALASNH TTEEFMARAISAIAELAKKAIEAIYRLADNHTTDKFMAAAIEAIALLATLAILAIALLASNHT TEEFMAKAISAIAELAKKAIEAIYRLADDHTSPTYIEKAIEAIEKIAKKAIKAIEMLAKNITTE EYQEKARKAILEILEKALEAIRRLEDNRR |

**Supplementary Table 8.** List of all designed two-component nanoparticles tested and their corresponding amino acid sequences including initiating methionines and $His_6$-tags. Designs that expressed solubly and co-eluted from IMAC are denoted in bold. Input oligomers from Supplementary Table 7 are included in parentheses.

| Antigen-fused component | Sequence |
| --- | --- |
| **BG505 SOSIP.v5.2(7S)–T33_dn2A without N241/N289** | **MKRGLCCVLLLCGAVFVSPSQEIHARFRRGARAENLWVTVY YGVPVWKDAETTLFCASDAKAYETKKHNVWATHCCVPTDP NPQEIHLENVTEEFNMWKNNMVEQMHTDIISLWDQSLKPCV KLTPLCVTLQCTNVTNNITDDMRGELKNCSFNMTTELRDKK QKVYSLFYRLDVVQINENQGNRSNNSNKEYRLINCNTSAITQ ACPKVSFEPIPIHYCAPAGFAILKCKDKKFNGTGPCPSVSTV QCTHGIKPVVSTQLLLNGSLAEEEVIIRSENITNNAKNILVQFN TPVQINCTRPNNNTVKSIRIGPGQWFYYTGDIIGDIRQAHCNV SKATWNETLGKVVKQLRKHFGNNTIIRFANSSGGDLEVTTH SFNCGGEFFYCNTSGLFNSTWISNTSVQGSNSTGSNDSITLP CRIKQIINMWQRIGQAMYAPPIQGVIRCVSNITGLILTRDGGST NSTTETFRPGGGDMRDNWRSELYKYKVVKIEPLGVAPTRCK RRVVGRRRRRRAVGIGAVSLGFLGAAGSTMGAASMTLTVQ ARNLLSGIVQQQSNLLRAPECQQHLLKDTHWGIKQLQARVL AVEHYLRDQQLLGIWGCSGKLICCTNVPWNSSWSNRNLSEI WDNMTWLQWDKEISNYTQIIYGLLEESQNQQEKNEQDLLEL DKWASLWGSMGNLAEKMYKAGNAMYRKGQYTIAIIAYTLA LLKDPNNAEAWYNLGNAAYKKGEYDEAIEAYQKALELDPN NAEAWYNLGNAYYKQGDYDEAIEYYKKALRLDPRNVDAIEN LIEAEEKQGAS** |
| **BG505 SOSIP.v5.2(7S)–T33_dn2A** | **MKRGLCCVLLLCGAVFVSPSQEIHARFRRGARAENLWVTVY YGVPVWKDAETTLFCASDAKAYETKKHNVWATHCCVPTDP NPQEIHLENVTEEFNMWKNNMVEQMHTDIISLWDQSLKPCV KLTPLCVTLQCTNVTNNITDDMRGELKNCSFNMTTELRDKK QKVYSLFYRLDVVQINENQGNRSNNSNKEYRLINCNTSAITQ ACPKVSFEPIPIHYCAPAGFAILKCKDKKFNGTGPCTNVSTV QCTHGIKPVVSTQLLLNGSLAEEEVIIRSENITNNAKNILVQLN ESVQINCTRPNNNTVKSIRIGPGQWFYYTGDIIGDIRQAHCNV SKATWNETLGKVVKQLRKHFGNNTIIRFANSSGGDLEVTTH SFNCGGEFFYCNTSGLFNSTWISNTSVQGSNSTGSNDSITLP CRIKQIINMWQRIGQAMYAPPIQGVIRCVSNITGLILTRDGGST NSTTETFRPGGGDMRDNWRSELYKYKVVKIEPLGVAPTRCK RRVVGRRRRRRAVGIGAVSLGFLGAAGSTMGAASMTLTVQ ARNLLSGIVQQQSNLLRAPECQQHLLKDTHWGIKQLQARVL AVEHYLRDQQLLGIWGCSGKLICCTNVPWNSSWSNRNLSEI WDNMTWLQWDKEISNYTQIIYGLLEESQNQQEKNEQDLLEL DKWASLWGSMGNLAEKMYKAGNAMYRKGQYTIAIIAYTLA LLKDPNNAEAWYNLGNAAYKKGEYDEAIEAYQKALELDPN NAEAWYNLGNAYYKQGDYDEAIEYYKKALRLDPRNVDAIEN LIEAEEKQGAS** |
| **BG505 SOSIP.v5.2(7S)–T33_dn10A** | **MKRGLCCVLLLCGAVFVSPSQEIHARFRRGARAENLWVTVY YGVPVWKDAETTLFCASDAKAYETKKHNVWATHCCVPTDP NPQEIHLENVTEEFNMWKNNMVEQMHTDIISLWDQSLKPCV KLTPLCVTLQCTNVTNNITDDMRGELKNCSFNMTTELRDKK QKVYSLFYRLDVVQINENQGNRSNNSNKEYRLINCNTSAITQ ACPKVSFEPIPIHYCAPAGFAILKCKDKKFNGTGPCTNVSTV QCTHGIKPVVSTQLLLNGSLAEEEVIIRSENITNNAKNILVQLN ESVQINCTRPNNNTVKSIRIGPGQWFYYTGDIIGDIRQAHCNV SKATWNETLGKVVKQLRKHFGNNTIIRFANSSGGDLEVTTH SFNCGGEFFYCNTSGLFNSTWISNTSVQGSNSTGSNDSITLP CRIKQIINMWQRIGQAMYAPPIQGVIRCVSNITGLILTRDGGST** |

| | |
|---|---|
| | **NSTTETFRPGGGDMRDNWRSELYKYKVVKIEPLGVAPTRCK RRVVGRRRRRRAVGIGAVSLGFLGAAGSTMGAASMTLTVQ ARNLLSGIVQQQSNLLRAPECQQHLLKDTHWGIKQLQARVL AVEHYLRDQQLLGIWGCSGKLICCTNVPWNSSWSNRNLSEI WDNMTWLQWDKEISNYTQIIYGLLEESQNQQEKNEQGSGS GSGSGGEEAELAYLLGELAYKLGEYRIAIRAYRIALKRDPNN AEAWYNLGNAYYKQGDYDEAIEYYQKALELDPNNAEAWYN LGNAYYKQGDYDEAIEYYEKALELDPENLEALQNLLNAMDK QG** |
| **BG505 SOSIP.v5.2(7S)–I53_dn5B** | **MKRGLCCVLLLCGAVFVSPSQEIHARFRRGARAENLWVTVY YGVPVWKDAETTLFCASDAKAYETKKHNVWATHCCVPTDP NPQEIHLENVTEEFNMWKNNMVEQMHTDIISLWDQSLKPCV KLTPLCVTLQCTNVTNNITDDMRGELKNCSFNMTTELRDKK QKVYSLFYRLDVVQINENQGNRSNNSNKEYRLINCNTSAITQ ACPKVSFEPIPIHYCAPAGFAILKCKDKKFNGTGPCTNVSTV QCTHGIKPVVSTQLLLNGSLAEEEVIIRSENITNNAKNILVQLN ESVQINCTRPNNNTVKSIRIGPGQWFYYTGDIIGDIRQAHCNV SKATWNETLGKVVKQLRKHFGNNTIIRFANSSGGDLEVTTH SFNCGGEFFYCNTSGLFNSTWISNTSVQGSNSTGSNDSITLP CRIKQIINMWQRIGQAMYAPPIQGVIRCVSNITGLILTRDGGST NSTTETFRPGGGDMRDNWRSELYKYKVVKIEPLGVAPTRCK RRVVGRRRRRRAVGIGAVSLGFLGAAGSTMGAASMTLTVQ ARNLLSGIVQQQSNLLRAPECQQHLLKDTHWGIKQLQARVL AVEHYLRDQQLLGIWGCSGKLICCTNVPWNSSWSNRNLSEI WDNMTWLQWDKEISNYTQIIYGLLEESQNQQEKNEQGSGS GSGSGGEEAELAYLLGELAYKLGEYRIAIRAYRIALKRDPNN AEAWYNLGNAYYKQGRYREAIEYYQKALELDPNNAEAWYN LGNAYYERGEYEEAIEYYRKALRLDPNNADAMQNLLNAKM REELEAS** |
| **HA–I53_dn5B** | **MKAILVVLLYTFTTANADTLCIGYHANNSTDTVDTVLEKNVTV THSVNLLEDKHNGKLCKLRGVAPLHLGKCNIAGWILGNPEC ESLSTASSWSYIVETSNSDNGTCFPGDFINYEELREQLSSVS SFERFEIFPKTSSWPNHDSNKGVTAACPHAGAKSFYKNLIW LVKKGNSYPKLNQSYINDKGKEVLVLWGIHHPSTTADQQSL YQNADAYVFVGTSRYSKKFKPEIATRPKVRDQEGRMNYYW TLVEPGDKITFEATGNLVVPRYAFTMERNAGSGIIISDTPVHD CNTTCQTPEGAINTSLPFQNIHPITIGKCPKYVKSTKLRLATG LRNVPSIQSRGLFGAIAGFIEGGWTGMVDGWYGYHHQNEQ GSGYAADLKSTQNAIDKITNKVNSVIEKMNTQFTAVGKEFNH LEKRIENLNKKVDDGFLDIWTYNAELLVLLENERTLDYHDSN VKNLYEKVRNQLKNNAKEIGNGCFEFYHKCDNTCMESVKN GTYDYPKYSEEAKLNREKIDGVSAEEAELAYLLGELAYKLG EYRIAIRAYRIALKRDPNNAEAWYNLGNAYYKQGRYREAIEY YQKALELDPNNAEAWYNLGNAYYERGEYEEAIEYYRKALRL DPNNADAMQNLLNAKMREEGGWELQHHHHHH** |
| **DS-Cav1–I53_dn5B** | **MELLILKANAITTILTAVTFCFASGQNITEEFYQSTCSAVSKG YLSALRTGWYTSVITIELSNIKENKCNGTDAKVKLIKQELDKY KNAVTELQLLMQSTPATNNRARRELPRFMNYTLNNAKKTN VTLSKKRKRRFLGFLLGVGSAIASGVAVCKVLHLEGEVNKIK SALLSTNKAVVSLSNGVSVLTFKVLDLKNYIDKQLLPILNKQ SCSISNIETVIEFQQKNNRLLEITREFSVNAGVTTPVSTYMLTN SELLSLINDMPITNDQKKLMSNNVQIVRQQSYSIMCIIKEEVL** |

| | |
|---|---|
| | **AYVVQLPLYGVIDTPCWKLHTSPLCTTNTKEGSNICLTRTDR<br>GWYCDNAGSVSFFPQAETCKVQSNRVFCDTMNSLTLPSEV<br>NLCNVDIFNPKYDCKIMTSKTDVSSSVITSLGAIVSCYGKTKC<br>TASNKNRGIIKTFSNGCDYVSNKGVDTVSVGNTLYYVNKQE<br>GKSLYVKGEPIINFYDPLVFPSDEFDASISQVNEKINQSLAFIR<br>KSDELLSAIGGSAEEAELAYLLGELAYKLGEYRIAIRAYRIAL<br>KRDPNNAEAWYNLGNAYYKQGRYREAIEYYQKALELDPNN<br>AEAWYNLGNAYYERGEYEEAIEYYRKALRLDPNNADAMQN<br>LLNAKMREEGGWELQHHHHHH** |

**Supplementary Table 9.** List of all antigen-fused components tested and their corresponding amino acid sequences.
